# Prenatal exposure to perfluoroalkyl substances modulates neonatal serum phospholipids, increasing risk of type 1 diabetes

**DOI:** 10.1101/588350

**Authors:** Aidan McGlinchey, Tim Sinioja, Santosh Lamichhane, Partho Sen, Johanna Bodin, Heli Siljander, Alex M. Dickens, Dawei Geng, Cecilia Carlsson, Daniel Duberg, Jorma Ilonen, Suvi M. Virtanen, Hubert Dirven, Hanne Friis Berntsen, Karin Zimmer, Unni C. Nygaard, Matej Orešič, Mikael Knip, Tuulia Hyötyläinen

**Author notes:** Equal contribution (Matej Orešič, Mikael Knip, Tuulia Hyötyläinen). Correspondence: Matej Orešič, Mikael Knip, Tuulia Hyötyläinen; MTM Research Centre, School of Science and Technology, Örebro University, 702 81 Örebro, Sweden.

## Abstract

In the last decade, increasing incidence of type 1 diabetes (T1D) stabilized in Finland, a phenomenon that coincides with tighter regulation of perfluoroalkyl substances (PFAS). Here, we quantified PFAS to examine their effects, during pregnancy, on lipid and immune-related markers of T1D risk in children. In a mother-infant cohort (264 dyads), high PFAS exposure during pregnancy associated with decreased cord serum phospholipids and progression to T1D-associated islet autoantibodies in the offspring. This PFAS-lipid association appears exacerbated by increased human leukocyte antigen-conferred risk of T1D in infants. Exposure to a single PFAS compound or a mixture of organic pollutants in non-obese diabetic mice resulted in a lipid profile characterized by a similar decrease in phospholipids, a marked increase of lithocholic acid, and accelerated insulitis. Our findings suggest that PFAS exposure during pregnancy contributes to risk and pathogenesis of T1D in offspring.

## Introduction

T1D is an autoimmune disease caused by destruction of insulin-secreting pancreatic beta-cells (Atkinson et al., 2014). The strongest genetic risk factors for T1D are found within the human leukocyte antigen (HLA) gene complex, yet only 3-10% of individuals carrying HLA-conferred disease susceptibility develop T1D (Achenbach et al., 2005). The role of environmental factors in T1D pathogenesis is thus obvious (Knip et al., 2005). We, and others, previously observed that children progressing to T1D-associated islet autoantibody positivity, or to overt T1D later in life, have a distinct lipidomic profile characterized by decreased blood phospholipid levels, including sphingomyelins (SMs), within the first months of life, preceding the onset of islet autoimmunity (Johnson et al., 2019; Orešič et al., 2008) and occurring even as early as at birth (Oresic et al., 2013). The cause of these metabolic changes is currently poorly understood. The gut microbiome is known to affect host lipid metabolism (Velagapudi et al., 2010) and is associated with progression to T1D (Kostic et al., 2015; Vatanen et al., 2018), particularly in the period after islet autoantibody seroconversion, but current data does not offer an explanation for the earlier changes in phospholipid levels (Kostic et al., 2015).

The incidence of T1D has been increasing over the last decades in many industrialized countries (Patterson et al., 2009). However, for unknown reasons, this has stabilized in the last decade, particularly in the Nordic countries (Harjutsalo et al., 2013). Environmental triggers and specific co-morbidities are often implicated in T1D, such as enterovirus infection, diet, and obesity (Knip et al., 2005), yet these factors do not explain this decrease in prevalence. Obesity, for example, has not shown a concomitant decrease since 2005 (Kaikkonen et al., 2012), and the number of severe enterovirus infections in Finland 2006-2010 increased, in fact, by 10 fold (Harjutsalo et al., 2013).

Notably, the time trend of human exposure levels to two widely-used industrial chemicals, namely, perfluorooctane sulfonate and perfluorooctanoic acid (PFOS and PFOA), does coincide with this trend in T1D incidence rate (Harjutsalo et al., 2013). These two compounds belong to the group of per- and poly-fluoroalkyl substances (PFAS) which are widely-used in food packaging materials, paper and textile coatings, and fire-fighting foams. The use of PFOS and PFOA has increased substantially since production started in the 1950s until the main, global manufacturer phased out its production of PFOS, PFOS-related substances and PFOA between 2000-2002. In the European Union, all uses of PFOS are now prohibited under Directive (2006/122/EC) which came into force in 2008 due to concerns regarding persistent effects in the environment and both the bioaccumulative and toxic effects in humans. PFOA is still manufactured, and a large number of other PFAS compounds remain in use. With a biological half-life of up to five years for PFOS and two to four years for PFOA in humans, concentrations of PFOS and PFOA started to decrease in man only after *ca*. 2005, with the levels of many other PFAS still showing increasing trends (Bjerregaard-Olesen et al., 2016). The main sources of exposure to PFAS in the general population are food and drinking water, with lesser sources including house dust and air. PFAS are transferred from mother to fetus *via* the placenta and to breast-fed infants *via* maternal milk (Croes et al., 2012).

Structurally, most PFAS resemble endogenous fatty acids, with fluorine substituted in place of hydrogen. Functionally, PFAS share some common features with bile acids, which are key metabolites involved in the digestion and absorption of lipids in the small intestine as well as in the maintenance of lipid and glucose homeostasis. Bile acids are excreted into the small intestine and subsequently reabsorbed, and similar enterohepatic circulation has been suggested for PFOS and PFOA (Zhao et al., 2015). It has been estimated that over 90% of PFOS and PFOA must be reabsorbed in order to explain the long half-life of these compounds in humans (Fujii et al., 2015). Bile acids can therefore potentially act as mediators, linking PFAS exposures and altered lipid metabolism.

There is a dearth of knowledge regarding PFAS as possible contributors to T1D risk / pathogenesis, although a contribution to the development of T1D has been proposed *via* impaired beta/immune-cell function and immunomodulation (Bodin et al., 2015). It has also been reported that PFOA and PFOS disrupt generation of human pancreatic progenitor cells (Liu et al., 2018). Prenatal and early-life exposure to perfluoroundecanoic acid (PFUnDA) aggravated insulitis development in NOD mice (Bodin et al., 2016). Recently, elevated levels of PFOS were reported in children at the point of diagnosis of T1D (Predieri et al., 2015).

Here we hypothesized that PFAS exposure *in utero* affects the phospholipid profile of newborn infants, which may contribute to increased T1D risk. In a mother-infant cohort study, we (i) analyzed metabolite profiles of pregnant mothers and their offspring at birth, (ii) quantified selected PFAS in maternal samples during pregnancy, and (iii) examined prenatal PFAS exposures in relation to neonatal metabolite profiles and progression to T1D-associated islet autoantibody positivity (AAb+) during follow-up. We then further experimentally examined the impact of PFAS exposure on both lipidomic and bile acid profiles as well as the development of insulitis / autoimmune diabetes in NOD mice, and finally verified our key findings in a prospective birth cohort study comprising children at risk for T1D.

## Results

### Metabolomic analyses of the mother-infant cohort

A total of 264 mother-infant dyads were included in the study (Fig. 1, Table S1). Maternal age at delivery was between 18.5 and 45.8 years, pre-pregnancy body mass index (BMI) was between 16.9 and 45.7 kg/m^2^, with 62% of the mothers being normal weight (BMI 18.5-25). All babies were born after gestational week 35. Seventy-four with HLA-conferred susceptibility for T1D were analyzed for AAb+ during follow-up, and ten among these progressed to at least one islet autoantibody.

**Fig. 1.**
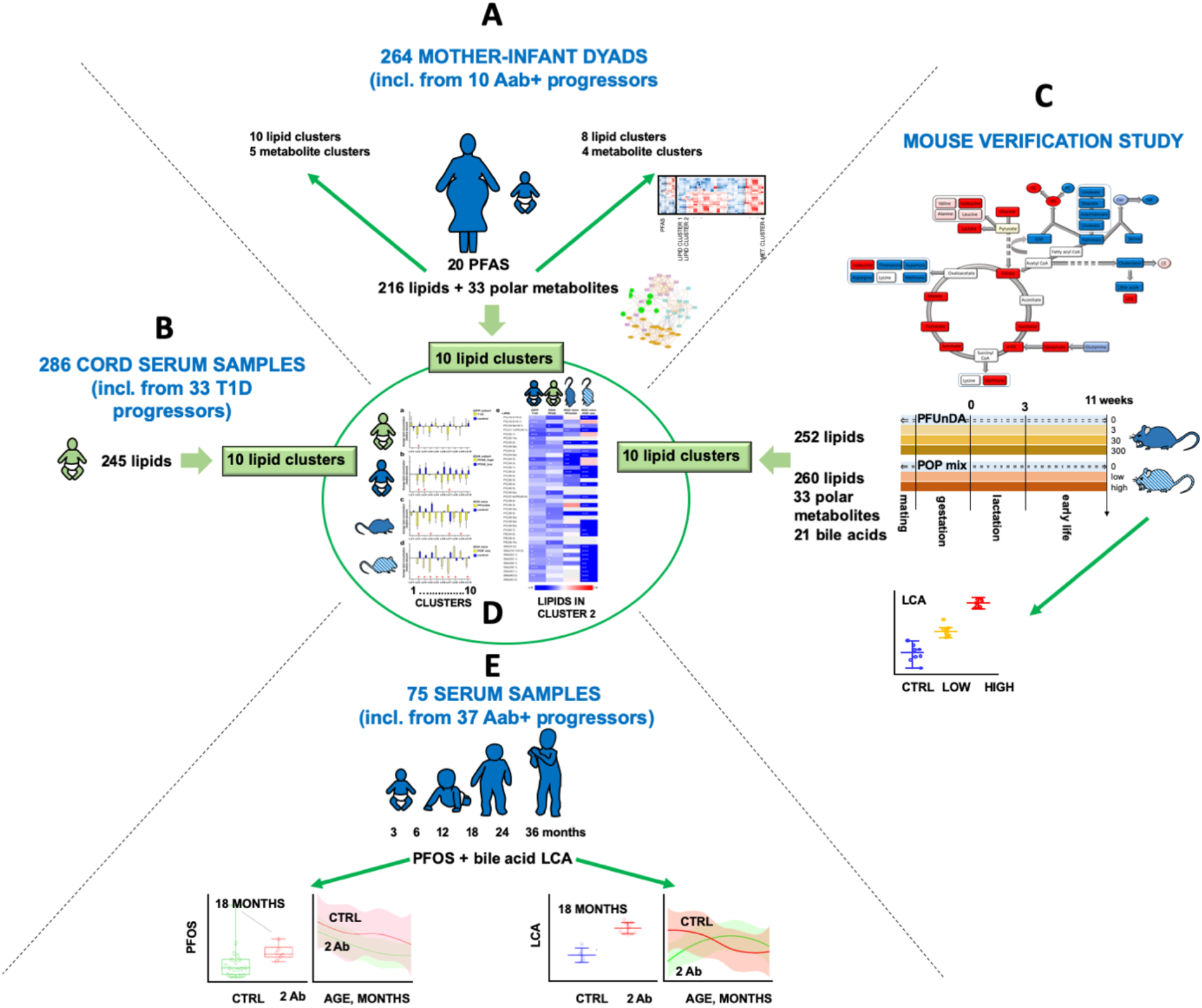
Overview of the workflow integrating prenatal PFAS exposure assessment, serum metabolomics and risk of type 1 diabetes. A. In the mother-infant cohort, PFAS levels and metabolomic profiles were determined from pregnant mothers, and metabolomics performed on cord serum from newborn infants. Metabolites were summarized as clusters, and associations between prenatal PFAS exposure and metabolomes were studied. B. Cord serum lipid changes due to prenatal PFAS exposure were then compared to previously-reported (Oresic et al., 2013) lipid-related differences between newborn infants who progressed to T1D later in life *vs.* those that remained healthy (Type 1 Diabetes Prediction and Prevention study – DIPP), and to (C) changes in lipid profiles brought on by exposure to a single PFAS compound or mixture of persistent organic pollutants, respectively, from two studies in non-obese diabetic (NOD) mice. D. The data across the four different studies (A, B, C) were summarized and compared by assigning lipids from each respective study to lipid clusters from the T1D study (Oresic et al., 2013). E. As a verification study, the exposure to PFOS and lithocholic acid levels were the further studied in the DIABIMMUNE prospective birth cohort.

Serum concentrations of 25 PFAS compounds were measured in the mothers during pregnancy, out of which 20 were detected (Table S2). The two most abundant PFAS were PFOS and PFOA, detected in all subjects. Our detected levels of PFOS and PFOA were lower than reported in previous studies (Bjerregaard-Olesen et al., 2016), most of which used samples collected before 2010, and therefore before recent, noted decreases in population blood levels of PFOS and PFOA (Land et al., 2018).

Metabolomic analyses were performed using two analytical platforms. Serum molecular lipids and polar metabolites were quantified from the mothers during pregnancy and from newborn infants (cord serum). Identified lipids (n = 216) and quantified polar metabolites (n = 34) were included in the final datasets (Data file S1). To reduce dimensional complexity and facilitate identification of global associations between metabolic profiles and maternal PFAS exposure, we first clustered the metabolites from all datasets into cluster variables using model-based clustering, followed by partial-correlation network analysis. The optimum number of clusters for each dataset, as assessed by Bayesian Information Criterion, returned ten Maternal Lipid Clusters (MLCs) and five Maternal polar Metabolite Clusters (MMCs), while the cord serum lipidomics data yielded eight Child Lipid Clusters (CLCs) and four Child polar Metabolite Clusters (CMCs) (Table S3).

### Metabolic profiles in mothers associate with PFAS exposure

PFAS exposure impacted the maternal metabolome (Fig. 2, Fig. S1). Total PFAS, as well as several individual PFAS levels were positively associated with MMC1 (amino acids, saturated free fatty acids and cholesterol). No strong associations were found between MLCs and individual PFAS exposures. For a subset of the cohort (n = 116), detailed lifestyle data, including dietary data during pregnancy, were available, and this dataset was used to estimate dietary sources of PFAS (Fig. S2). Shellfish showed the strongest correlation with serum PFAS levels, with other food items also showing significant associations with PFAS levels, such as fish, cereals and fruit juice. This suggests that, to a large extent, seafood consumption drives maternal PFAS levels.

**Fig. 2.**
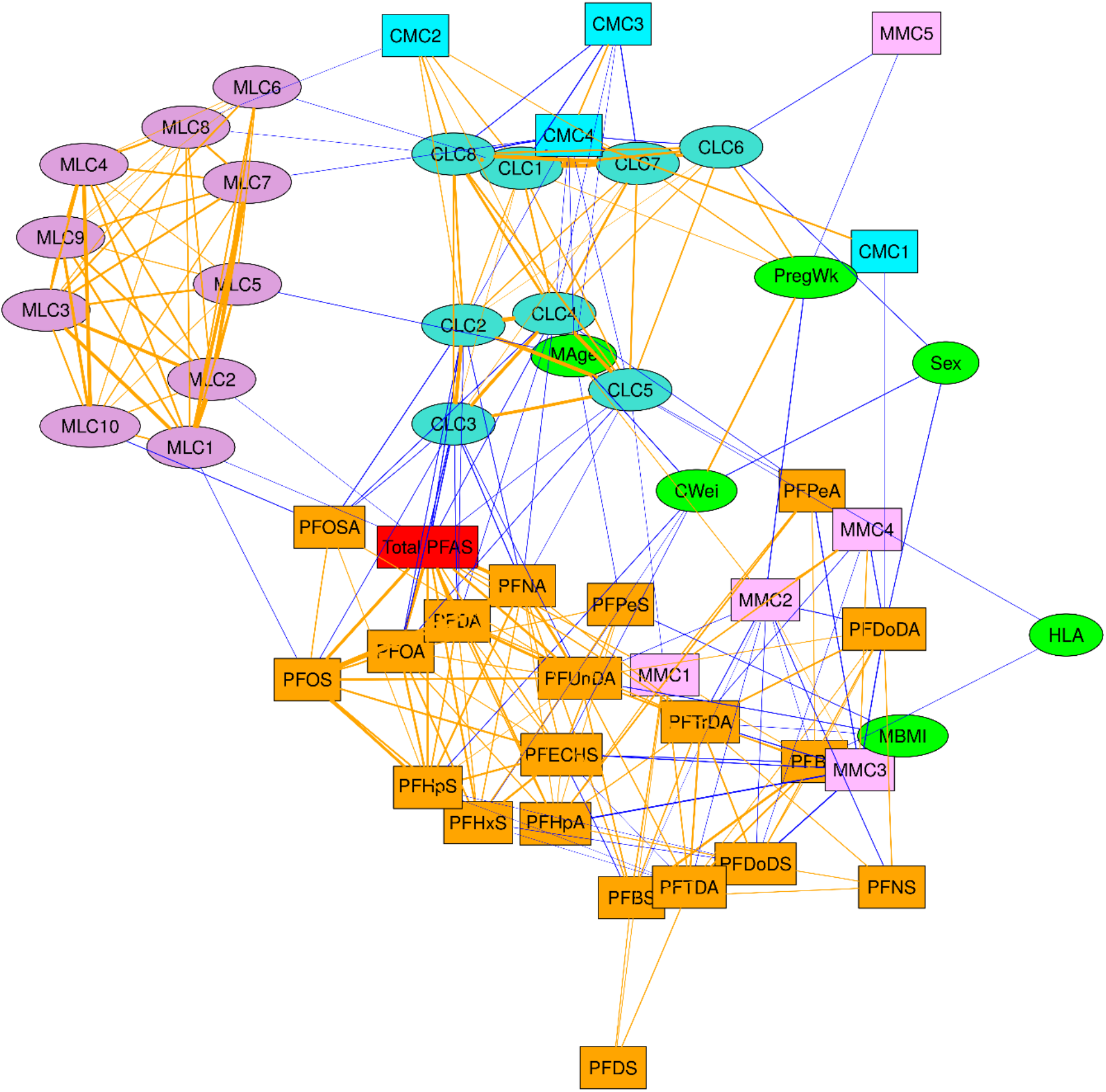
Partial correlation network showing associations between demographic data, maternal PFAS levels and lipidome / metabolome cluster variables from mothers and their newborn infants. The network was constructed using the *qpgraph* R package. Node color / shape represents the different datasets, edge color denotes a positive (orange) or inverse (blue) association. The threshold non-rejection rate was set as 0.4. Node abbreviations: PregWk, weeks of pregnancy; HLA, HLA risk locus (1 = lower risk, 2 = higher risk); CWei, child birth weight; MBMI, maternal BMI; MAge, maternal age.

In agreement with prior findings (Tsai et al., 2018), we observed inverse correlations between the number of the mothers’ previous deliveries and levels of specific PFAS: PFOA (r = −0.44, FDR q < 0.01), PFHpS (r = −0.39, q < 0.01), PFOS (r = −0.31, q = 0.07), total PFAS (r = – 0.26, q = 0.23, nominal p = 0.006).

### Cord serum metabolic profiles in newborn infants associate with PFAS exposure

Partial correlation network analysis revealed a marked impact of maternal PFAS exposure on the cord serum metabolome of newborn infants (Fig. 2). Inverse associations between cord serum lipids and PFOS, PFOA and total PFAS exposure were observed, particularly for clusters CLC2 (sphingomyelins (SMs), abundant phosphatidylcholines (PCs)), CLC3 (lysophosphatidylcholines (LPCs)), and CLC4 (PUFA-containing phosphatidylcholines PCs). CMC4 (mainly specific amino acids) was positively associated with PFOS and perfluorodecanoic acid (PFDA) exposure.

Next, the infants were classified into four groups (quartiles) based on total maternal PFAS exposure levels, as a sum of all individual PFAS levels. Among the eight CLCs and four CMCs, two lipid clusters (CLC2: ANOVA p=0.035, CLC3: Tukey’s HSD p=0.0062) and one polar metabolite cluster (CMC4: Tukey’s HSD p=0.0035). These Tukey’s HSDs show differences between the highest (Q4) and lowest (Q1) PFAS exposure quartiles.

Linear regression (LR) with regularization was performed to determine the relative contribution of individual maternal PFAS on CLC2, CLC3 and CMC4. LR modelling showed that, maternal PFAS such as: PFOSA, PFPeA, PFOA, PFNA were the top linear predictors (ridge regression coefficients (s) ~ 0.8) of the cord serum lipid profiles in CLC2 (SMs, abundant PCs) and CLC3 (LPCs) (Fig S3A-B.). Moreover, it suggested that maternal exposure to PFHxS, PFHpS, PFUnDA and PFNS might affect cord serum lipids, however, the strength of these associations was comparatively weaker. Regression of PFAS to CMC4 (amino acid cluster) identified PFPeA, PFDA, PFOS as the top linear predictors (s > 0.8) of cord serum amino acids grouped in CMC4.

At the individual metabolite level, (i) 39 molecular lipids, mainly decreased LPCs, SMs and PCs (from CLC2, CLC3, and to a lesser extent, CLC4), and (ii) 10 polar metabolites (increased amino acids from CMC1 and, mainly, CMC4), were altered between higher and lower quartiles of total maternal PFAS exposure (p < 0.05, ANOVA Tukey HSD). In order to access the effect of individual maternal PFAS on cord serum lipid and polar metabolite levels, we performed LR with the maternal PFAS as predictors of selected representative cord serum metabolite concentrations. Metabolites that were significantly altered between higher and lower quartiles of total maternal PFAS exposure were considered for LR modelling. Regression analysis showed that, PFNA, PFOS, PFDA, were the top linear predictors (s > 0.7) of cord serum LPC(20:4) (from CLC3) (Fig. 3A-C), whilst PFNA, PFPeA, PFOA, were the linear predictors (s > 0.5) of SM(d38:1) (from CLC2) (Fig. 3D-F). PFHxS was identified as a top predictor of cord serum methionine (from CMC4) levels, along with PFDA, PFOS, PFNA (Fig. 3G-I).

**Fig. 3.**
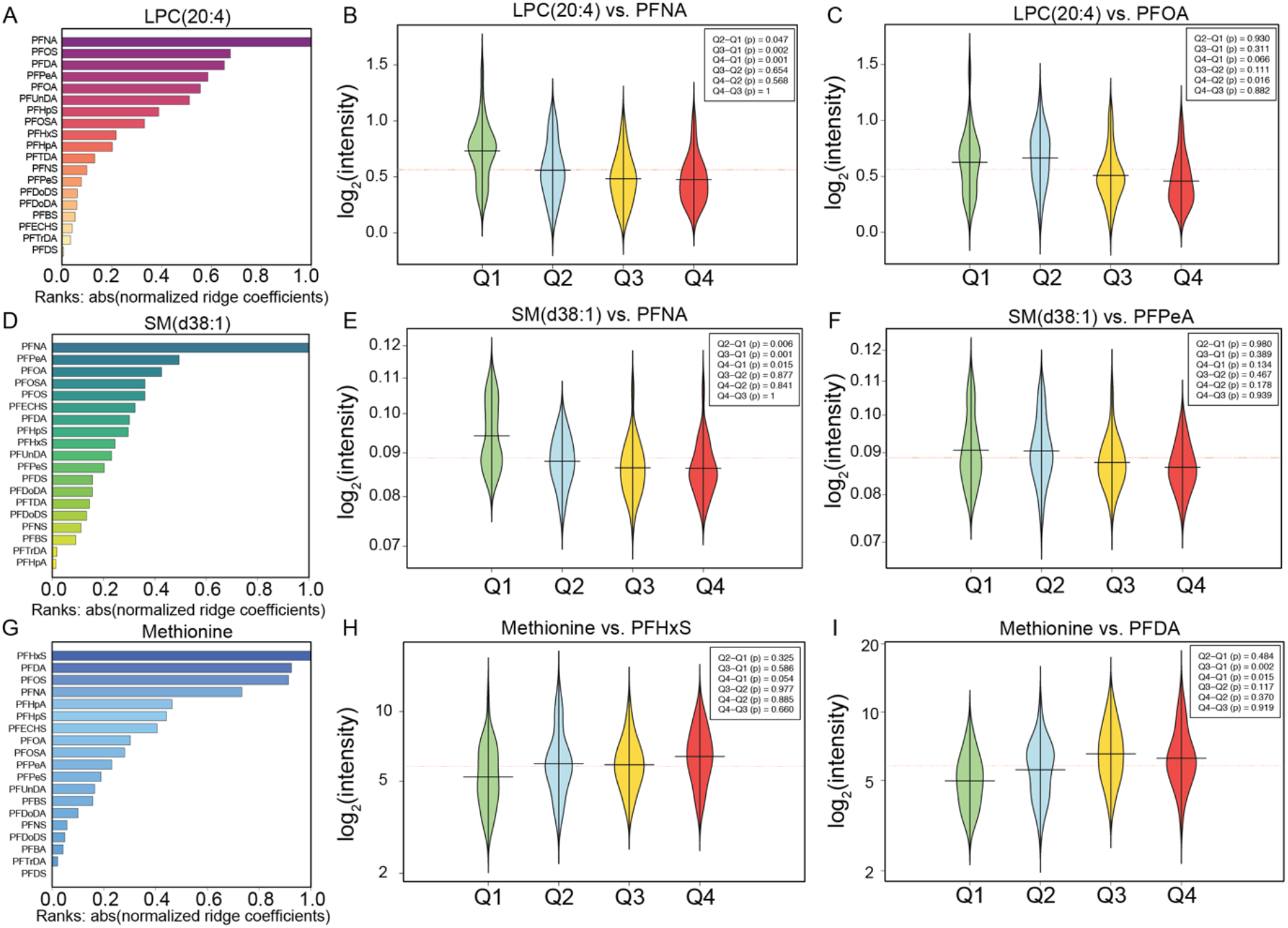
Impact of prenatal exposure of selected PFAS on specific lipids. A, D, G) Horizontal bar plots showing ranks of different PFAS as predictors of LPC(20:4), SM(d38:1) and Methionine levels, respectively, as measured in cord serum. These metabolites are significantly (adjusted p < 0.05) altered between highest (Q4) and the lowest (Q1) quartiles of total maternal PFAS levels. The PFAS are ranked and sorted by their absolute normalized regression (ridge) coefficients, *i.e.* the PFAS that has a higher impact on the metabolic profiles are shown at the top of the chart. B-C, E-F, & H-I) Beanplots showing levels of LPC(20:4), SM(d38:1) and Methionine, measured in cord serum, which are associated with the top two contributing maternal PFAS. The intensities of metabolites are plotted across different quartiles (Q1– Q4) of the contributing maternal PFAS. Red, horizontal bars indicate population mean, black horizontal bars are group mean, and “bean” width represents the density of samples in a group/quartile. The mean difference in the metabolite intensities along different quartiles (Qs) were compared by ANOVA and post-hoc Tukeys’ HSD test.

### Impact of PFAS exposure on cord serum lipids associated with risk of T1D progression

As the cord serum lipid profile associated with total maternal PFAS exposure here proved similar to that found previously as being associated with progression to T1D (Oresic et al., 2013), we also examined the impact of PFAS exposure on T1D-associated lipids. First, we assigned the lipids from the present study to the same lipid clusters (LCs) as used in our previous study, and investigated their association with PFAS levels.

Of the ten lipid clusters used in the earlier study, four showed significant differences between the highest and lowest exposure groups: LC2 (ANOVA Tukey’s HSD test, p = 0.003), LC3 (p=0.001), LC6 (p=0.011) and LC7 (p=0.001). In our previous study, the most significantly-changing lipid clusters associated with T1D progression were LC2 (major PCs) and LC7 (SMs), which were down-regulated in newborn infants who later progressed to clinical T1D. In agreement with these results, the lipid levels from the present study in those same clusters were also reduced in the highest (Q4) PFAS exposure group by comparison to the lowest (Q1) exposure levels in the current study. In addition, lipid clusters LC3 (LPCs) and LC6 (PUFA-containing phospholipids) showed clear differences in the current study, with lower lipid levels again seen in the highest exposure group. Likewise, to identify which maternal PFAS are the key contributors to these mean lipid changes in clusters LC2 and LC7, we performed LR modelling (Fig. S3 D-F). Regression analysis showed that PFHxS and PFDA were the top linear predictors (s > 0.8) of the lipids grouped in cluster LC2 (major PCs) and LC7 (SMs).

Given the observed impact of prenatal exposure to PFAS on cord serum lipids associated with progression to T1D, we also examined whether HLA-conferred risk of T1D plays a role in mediating the impact of PFAS exposure on lipids in newborn infants. We divided the infants into two categories according to HLA-associated T1D risk: (low *vs.* increased; Table S1) and two categories according to prenatal total PFAS exposure (quartiles 1 & 2 *vs*. 3 & 4). A multi-way ANOVA was then performed across these groups for the 39 lipids found associated with PFAS exposure. When examining the interaction effect between HLA risk and PFAS exposure, we found eight lipids with a p-value < 0.05 (Table S4).

In a subset of the cohort, which included 74 children enrolled in the clinical trial and for whom follow-up data were available, we then compared prenatal PFAS exposure in children who progressed to islet autoantibody positivity during the follow-up (n = 10) *vs*. those that did not. We found that the children who progressed to at least one islet autoantibody had elevated prenatal levels of several PFAS. The total maternal PFAS levels of the children who seroconverted to at least autoantibody (AAb+) were significantly different (two-sample t-test, p=0.035) from the serum antibody negative group (AAb-, control).

In order to estimate the ranks of individual PFAS (predictors) for the separation of the AAb+ *vs*. AAb-groups, we fitted a logistic ridge regression (LRR) model (mean AUC = 0.78, 95% CI: 0.51 – 0.76) with 20 PFAS and estimated their ranks (absolute difference in odds ratios from unity) based on their impact on classifying the AAb+ *vs.* AAb-groups. The LRR model showed that at least five PFAS were linear predictors of antibody positivity, with PFHxS and PFHpS having the highest odds ratios (Fig. 4A). Next, we determined the effect of different PFAS, either singly or in combination, on the separation of AAb+ vs. AAb-groups, by developing stepwise LRR models. This LRR modelling identified an optimal set of five PFAS that aided in the separation of AAb+ vs. AAb-groups at (AUC = 0.81, 95% CI: 0.76 – 0.85) (Fig. 4B).

**Fig. 4.**
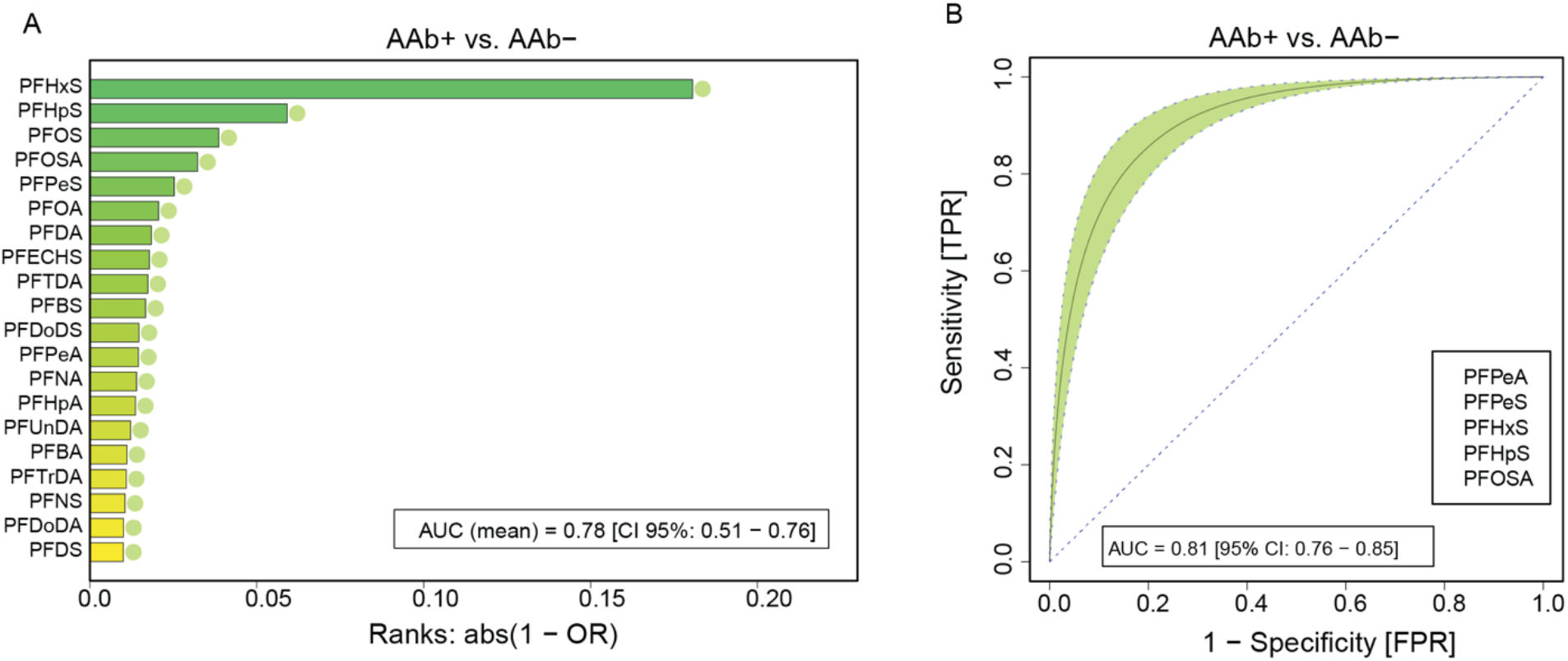
Logistic ridge regression (LRR) models showing PFAS exposure as linear predictors of autoantibody AAb+ vs. AAb-, in a follow-up EDIA study. (A) Ranks (abs (1-OR)) of predictors (PFAS) obtained from the LRR model, adjusted by mother’s age and BMI are shown. ‘OR’ depicts odds ratios. The most contributing PFAS (predictor) that aided in the separation of AAb+ *vs.* AAb- (mean AUC = 0.78, 95% CI: 0.51 – 0.76) are shown in the top of the chart. (B) shows the receiver operating characteristic (ROC) from stepwise – predictive LRR model (10-fold cross-validation). An optimal set of five PFAS (AUC = 0.81, 95% CI: 0.76 – 0.85) were associated with AA+ *vs.* AAb-. The green-shaded region depicts the 95% confidence region.

### Pre- and postnatal PFAS exposure in NOD mice alters offspring lipid profiles

Based on the metabolomics results from the mother-infant cohort, we hypothesized that PFAS exposure during pregnancy has a contributing, causal impact on phospholipid levels, which, in turn, associates with increased risk of T1D. Two previous studies in NOD mice suggest that maternal PFAS exposure accelerates insulitis development and progression to autoimmune diabetes (Berntsen et al., 2018; Bodin et al., 2016). We analyzed serum lipidomic profiles from these two studies which exposed 11-week-old NOD mice to either (i) a mixture of persistent organic pollutants (POPs) in feed (including a total PFAS intake 0 (control), 0.14 (low), or 2.8μg/day (high)). Here the low level corresponds to the approximate level of PFAS in human serum and whilst the high level representing a level 50-times higher than the level of total PFAS in serum (Berntsen et al., 2017) (Data file S2), or to (ii) PFUnDA at varying levels in drinking water (0, 3, 30 and 300 μg/L) (Data file S3) (Bodin et al., 2016). Additionally, analyses of polar metabolites and bile acids were performed in the POP mixture exposure study. It should be noted that the levels of the other (non-PFAS) POPs were significantly lower than those of PFAS (Berntsen et al., 2017). These exposures commenced at the times of mating, during gestation and lactation and until 11 weeks of age (Berntsen et al., 2018).

Marked changes were observed in the POP study, with the strongest effects seen in the high exposure group, but with significant changes occurring also in the low exposure group, corresponding to expected human exposure levels. A total of 143 out of 183 identified lipid levels, along with the levels of 16 polar metabolites out of 65, were significantly changed. Specifically, exposure caused a marked reduction in the levels of a large number of phospholipids, with several PUFA-containing TGs being significantly down-regulated as well. We also identified significant changes in the levels of several free fatty acids, free cholesterol, amino acids, glycerol-3-phosphate and 3-hydroxybutyric acid, particularly in the high exposure group, with significant upregulation of TCA cycle metabolites.

Exposure to only a single PFAS, PFUnDA, also caused significant changes in lipid profiles at the highest exposure level, with similar patterns of changes found with the two lower concentrations, although these did not reach statistical significance. A total of 32 out of 126 identified lipids were significantly different between the high exposure and unexposed groups, with the changes mainly due to decreased levels of PCs and SMs.

Next, we assigned the measured lipids to the same lipid clusters (LCs) as in our previous study (Oresic et al., 2013) and investigated the association of PFAS exposure with these known lipid clusters (Fig. 5). In mice exposed to the POP mixture, eight of ten clusters showed significant changes between control and high exposure groups, and two clusters changed significantly between control and low exposure groups. In agreement with our previous study and our mother-infant cohort study presented here, LC2 decreased significantly with increasing PFAS exposure. In addition, clusters LC4, LC9 and LC10 showed marked differences in the current study with significantly decreased lipid levels in the highest exposure group. These clusters contained mainly minor phospholipids, major TGs and long-chain PUFA-containing TGs. Of the ten lipid clusters, four showed significant differences between the highest PFUnDA-exposure group and the control mice. One lipid cluster showed a significant difference even at the lowest level of exposure compared to control.

**Fig. 5.**
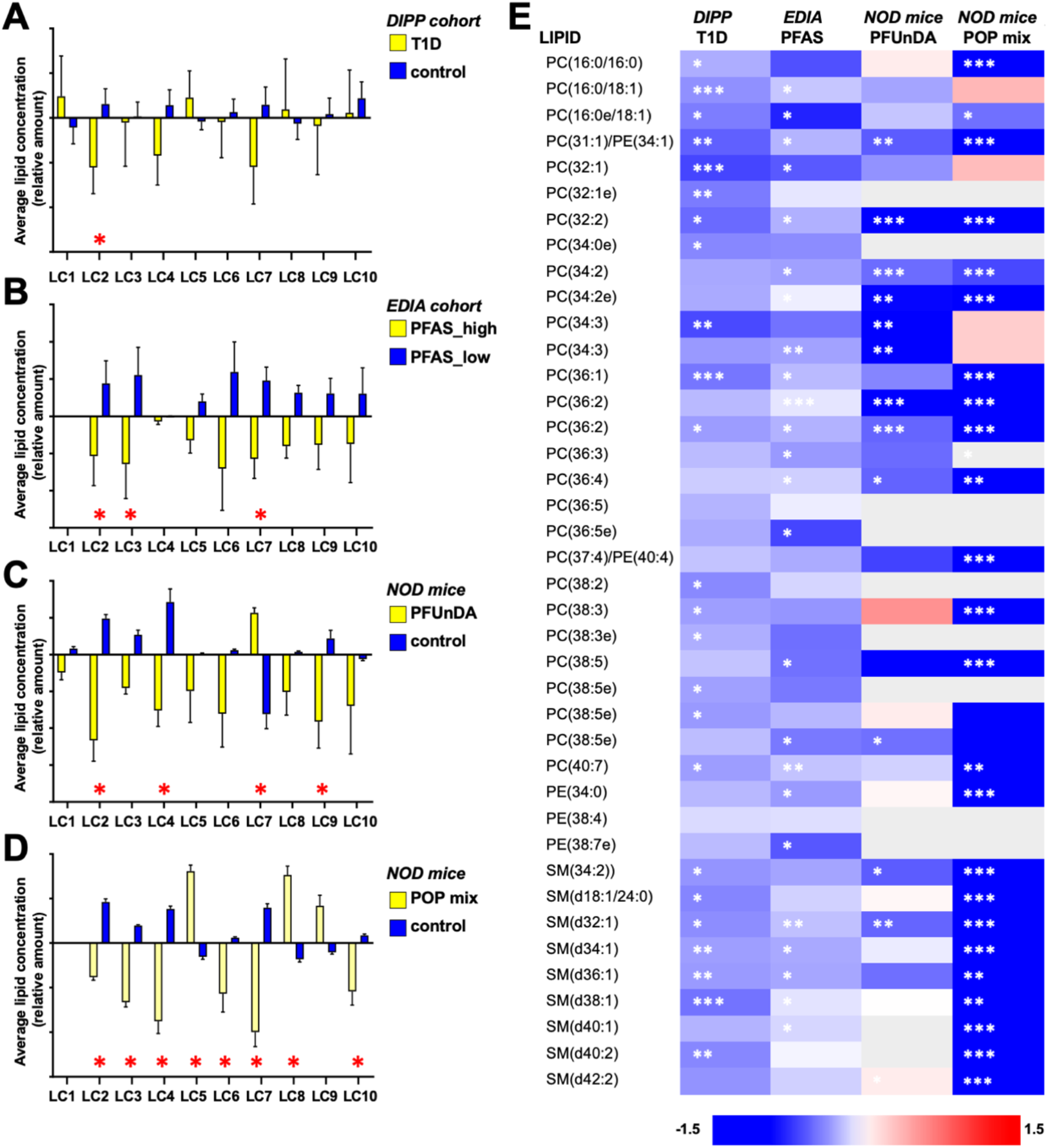
Comparison of lipidomic profiles across four different studies, using lipid cluster assignments from an earlier study (Oresic et al., 2013). A. Cord serum profiles from progressors to T1D (yellow bars) and control children (blue bars), from a previous report in the Diabetes Prediction and Prevention (DIPP) study in Finland (Oresic et al., 2013). B. Cord serum from mother-infant (EDIA) cohort, with high PFAS exposure (yellow) and low exposure (blue). C. NOD mice exposed to a high level of PFUnDA (yellow) and unexposed control mice (blue). D. NOD mice exposed to POP mixture at a high level (yellow) and unexposed mice (blue). E. Fold changes between the groups in A-D of lipids in cluster LC2. Statistical significance levels: *p<0.05, **p<0.01, ***p<0.001.

Using the cluster assignments from the earlier study (Oresic et al., 2013), a remarkable similarity was observed when comparing the results across all four studies (Fig. 5): (i) a previously-reported study of the cord serum lipidome in relation to progression to T1D (Oresic et al., 2013), (ii) the association between PFAS exposure and cord serum lipid profiles in the mother-infant cohort presented here, and (iii) the effects of PFUnDA and (iv) a PFAS-containing POP mixture on the lipid profiles of NOD mice. Among the 15 individual lipids reported to have significant association with the development of T1D, 11 and 10 of these were detected also in the NOD mice, exposed to PFUnDA and the POP mixture, respectively. Among these lipids, three showed significant differences between control and high exposure groups in the PFUnDA model and nine in the POP mixture exposure model. Notably, the pattern of the changes in the two mouse models were very similar, and *in vitro* exposure of macrophages to groups of chemicals from the POP mixture revealed that the PFAS mixture was driving the differences also in the POP mixture model (Berntsen et al., 2018; Bodin et al., 2016). Strikingly, all lipid changes taking place in association with higher PFAS exposure occurred in the same direction as reported previously in relation to increased risk of T1D.

### High exposure to PFAS associates with elevated levels of lithocholic acid and islet autoantibody positivity

As bile acids and PFAS utilize similar enterohepatic circulation (Zhao et al., 2015), we also examined the impact of the POP mixture on serum bile acid levels in serum of NOD mice. Indeed, the bile acid profiles were markedly altered, in a dose-dependent manner, on exposure to the POP mixture (Table S5). A majority of the bile acids, including the primary bile acids (CA, CDCA) were downregulated, while lithocholic acid (LCA) was markedly upregulated in comparison to the control group (Fig. 6A; fold changes of 2.1 and 5.9 at low and high exposure to POP, p-values of 6.7×10^-4^ and 5.6×10^-8^, respectively). Notably, there was a strong inverse association between the levels of LCA and the levels of SMs and LPCs (two examples shown in Fig. 6B-C). Specifically, all SMs were downregulated (median R = −0.63, p = 0.000071-0.04) while 70% of the LPCs were significantly inversely associated with the LCA (median R = −0.60, p = 4.1×10^-7^-0.01), except for LPC(22:3), which was positively correlated with LCA (R=0.54, p=0.007).

**Fig. 6.**
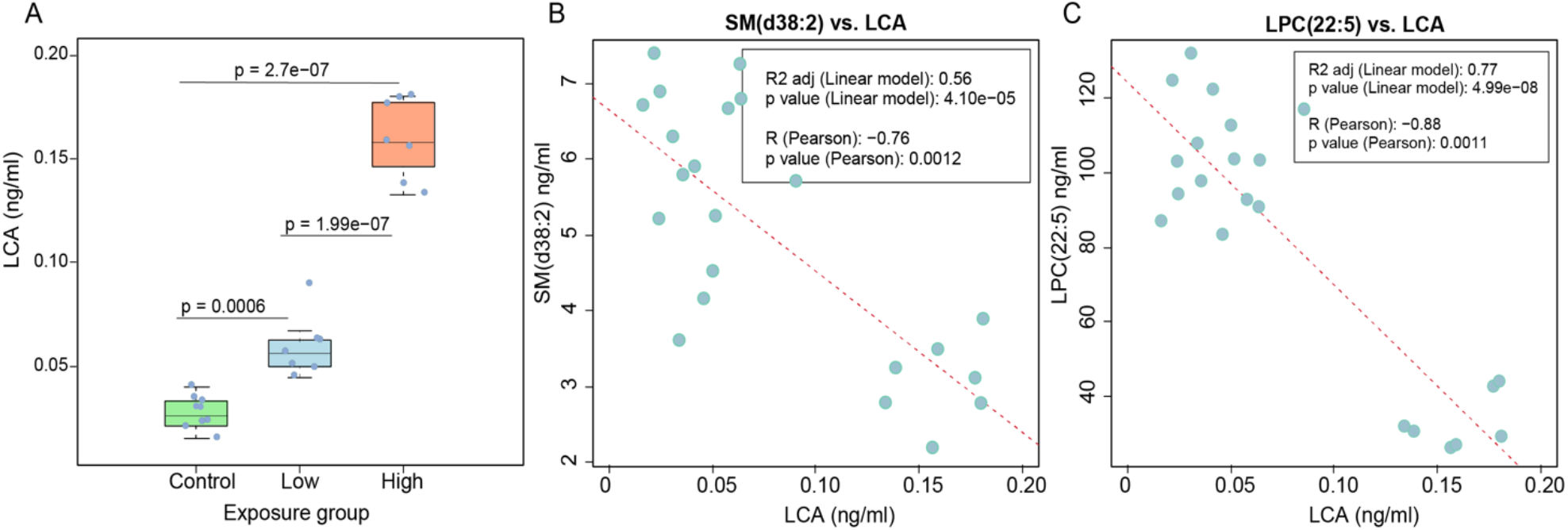
Impact of POP mixture on levels of lithocholic acid (LCA) in NOD mice. A. LCA levels in NOD mice exposed to POP mixture at two different doses. Associations of LCA levels in NOD mice exposed to POP mixture, with B. SM(d38:2) and C. LPC(22:5).

Next, we compared the plasma PFOS and lithocholic acid (LCA) concentration differences between the children who progressed to multiple islet autoantibodies (mAAb+) *vs.* controls (CTR), who remained AAb negative, in a previously-reported subgroup of the DIABIMMUNE study (Kostic et al., 2015) (Data file S4). In line with our findings in the mother-child cohort, we observed a higher level of PFOS, particularly in breast-fed (≥ 30 days) children who progressed to mAAb+ (n = 6) than in CTR (n = 20) at 18 months of age (p-value < 0.05, Fig. S4A). In addition, we found that in the longitudinal profile, PFOS remained persistently higher in the mAAb+ group than in CTR (Fig. S4B). We then sought to determine whether LCA levels were altered with exposure. We found that children with the highest level (Q4) of PFOS exposure tended to have increased levels of LCA compared with children who had a low level (Q1) of PFOS exposure (Fig. S4C). We also observed that LCA differed between the cases and controls. Higher levels of LCA were found in children who progressed to mAAb+ than in the CTR at 6 and 36 months of age (p-value < 0.05, Fig. S4D-F).

## Discussion

By integrating PFAS exposure and metabolomic data from pregnant mothers with metabolomic data from their newborn infants, we were able to demonstrate altered cord serum metabolic signatures associated with high PFAS exposure during pregnancy and subsequently verify these findings in NOD mouse models of pre- and postnatal PFAS exposure. We also reported a remarkable similarity between the metabolic signature observed in the current (EDIA) study and the known signature associated with progression to T1D.

The composition of the cord blood metabolome reflects maternal metabolism, placental transfer across the maternal-fetal axis as well as fetal metabolism itself (Hart et al., 2008). This may explain the weak associations between metabolic profiles of mothers and their offspring. The observed PFAS-associated metabolic changes seen in cord blood were not associated with PFAS-related maternal metabolic changes. These fetal metabolic changes are therefore likely the result of PFAS exposure itself, rather than a downstream consequence of maternal metabolic changes. Several studies have shown that maternal levels of PFAS are reflected in the developing fetus (Winkens et al., 2017) and there is a strong correlation between PFAS levels in maternal and cord blood (Kim et al., 2011). One recent study indicates that PFAS concentrations in first trimester fetuses represent 5% to 27% of maternal plasma concentrations, fetal concentrations increasing with gestational age (Mamsen et al., 2017). A comparison of transplacental transfer efficiency for different PFAS suggests an inverse relationship with the chain-length of the perfluoroalkyl group and a somewhat lower transfer efficiency for perfluorosulfonic acids compared to perfluorocarboxylic acids (Winkens et al., 2017). We did not determine PFAS levels in newborn infants due to the limited volumes of samples available for quantification.

There is general consensus that exposure to PFOA and PFOS alters the immune system in experimental models, with documented effects including altered antibody and cytokine production (DeWitt et al., 2009). In our study, we observed that prenatal PFAS exposure caused decreased levels of several phospholipids, particularly SMs and specific PCs, which were previously found to be persistently down-regulated in children who later progressed to islet autoimmunity (Johnson et al., 2019) and clinical T1D (Orešič et al., 2008). The importance of sphingolipid metabolism in the pathogenesis of T1D was highlighted by a genome-wide association study which identified eight gene polymorphisms involved in sphingolipid metabolism which contribute to T1D predisposition, and levels of which also correlated with the degree of islet autoimmunity in patients with recent-onset T1D (Holm et al., 2018). Recently, altered sphingolipid metabolism was also observed in peripheral blood mononuclear cells (PBMCs) from children who later went on to develop overt T1D (Sen et al., 2020). Among the PFAS measured in our study, the main implicated drivers of the observed changes in cord serum phospholipid levels were PFOS, PFOSA, PFOA, PFDA, and PFNA. Also, serine and palmitic acid (precursors of SMs) were found to be down-regulated with higher PFAS exposure and correlated with SM levels (R > 0.4), both in newborn infants as well as in NOD mice, where the exposure to PFAS was also associated with accelerated insulitis development (Bodin et al., 2016). We conclude that high PFAS exposure may alter sphingolipid levels during fetal development which might then go on to play a pathogenic role in the development of T1D later in life. The potential role of HLA-associated T1D risk in exacerbating the effect of prenatal PFAS exposure on lipid levels in the offspring, as suggested by our data, clearly demands further investigation.

Altered bile acid levels as observed in NOD mice exposed to POP mixtures, and in children positive for multiple islet autoantibodies, may explain altered lipid profiles. In animal models, LCA exposure has been shown to cause downregulation of circulating LPCs and SMs (Matsubara et al., 2011), which is precisely what we have observed both in previous T1D studies (linking early lipid changes with progression to T1D later in life) (Orešič et al., 2008) as well as in the current study in relation to PFAS exposure. Notably, LCA and its metabolites were recently found to control host immune responses by modulating the balance of TH17 and Treg cells (Hang et al., 2019). LCA is also an agonist for membrane receptor TGR5 which mediates the release of glucagon-like peptide 1 (GLP-1), promoting insulin release from pancreatic beta-cells (Kumar et al., 2012). Bile acid metabolism is closely linked with gut microbial activity and, indeed, PFAS exposure has been shown to cause reduced microbiome diversity in infants (Iszatt et al., 2019). In the DIABIMMUNE cohort, including the samples studied here, the children that progressed to multiple islet autoantibodies later in life were found to have decreased alpha-diversity in their gut microbiota. Decreased levels of LCA and increased levels of SMs in stool were also associated with relative overabundance of pathobionts, including AAb+-associated *Ruminococcus* (Kostic et al., 2015). Our data presented in the current study thus suggests that PFAS impact absorption of bile acids, which may, in turn, affect circulating lipid levels and immune system homeostasis.

We also observed an inverse association between maternal levels of specific PFAS and the number of previous deliveries, due to transfer of PFAS to the fetus, and excretion *via* breast milk (*i.e.*, breast feeding) which is in line with earlier reports (Tsai et al., 2018). Interestingly, pooled analysis across multiple studies suggests that increasing birth order is associated with lower risk of T1D (Cardwell et al., 2011). Our findings therefore support the notion that decreased maternal PFAS levels, due to multiple pregnancies, is one possible cause of this previously unexplained phenomenon.

Taken together, we conclude that high prenatal exposure to PFAS appears to alter lipid profiles in newborn infants, which, in turn, may increase the risk of islet autoimmunity and T1D. Our data also highlight a potential role for a gene-environment interaction (HLA risk genotype and prenatal PFAS exposure), which may lead to altered lipid profiles in newborn infants at-risk of developing T1D. Our findings may offer an explanation for the changing trend in the incidence of T1D in certain Western countries as well as underscore the need for investigation of how exposure to specific PFAS and other persistent chemical pollutants during pregnancy and early childhood affect the risk and pathogenesis of T1D.

## Materials and Methods

### Experimental design

The study setting is shown in Fig. 1. Associations between prenatal PFAS exposure and metabolomes were first studied in the mother-child cohort (EDIA study). PFAS levels and metabolomic profiles were determined from pregnant mothers, and metabolomics performed on cord serum from newborn infants. Next, changes in lipid profiles brought on by exposure to a single PFAS compound or mixture of persistent organic pollutants, respectively, were examined from two studies in non-obese diabetic (NOD) mice. We also compared cord serum lipid changes due to prenatal PFAS exposure from the EDIA study to previously-reported, lipid-related differences between newborn infants who progressed to T1D later in life *vs.* those that remained healthy (Oresic et al., 2013). We then compared the similarities of lipid profiles across the four studies. Finally, the key findings from the study were verified in the DIABIMMUNE birth cohort.

### Mother-infant cohort

Pregnant women were recruited from January 28, 2013 to February 26, 2015, in the context of the EDIA (Early Dietary Intervention and Later Signs of Beta-Cell Autoimmunity: Potential Mechanisms) study, which is a small-scale intervention trial comparing weaning infants onto an extensively-hydrolyzed milk formula *vs*. a conventional cow’s milk-based formula. Families were contacted at the time of the fetal ultrasonography visit, which is arranged for all pregnant women in Finland around gestational week 20. Written, informed consent was signed by the parents to permit analysis of their HLA genotype to exclude infants without HLA-conferred susceptibility to T1D. At this point, 68% of the infants to be born were excluded. Separate informed consent was obtained from eligible parents at the beginning of the third trimester to analyze the offspring’s genotype and to continue in the intervention study.

The cord blood from 309 newborn infants was screened to determine the HLA genotype, as previously described (Hermann et al., 2003). The degree of HLA susceptibility to T1D was divided into six categories (high-risk, moderate-risk, low-risk, neutral, protective and strongly protective genotypes), as earlier defined (Ilonen et al., 2016). A total of 89 infants were eligible for participation in the intervention study, carrying high-risk and moderate-risk genotypes. In that study, 82 infants were randomized and 73 remained in follow-up until the age of 12 months.

For the current study, the HLA risk categories were combined into two classes; the increased risk genotypes and the low-risk genotypes. Genotypes where HLA-(DR3)-DQA1*05-DQB*02 and/or DRB1*04:01/2/4/5-DQA1*03-DQB1*03:02 were present with each other, homozygous or heterozygous with a neutral haplotype were classified as increased risk and all other genotypes as low risk. Maternal diet during pregnancy was assessed by validated, semiquantitative food frequency questionnaire (Erkkola et al., 2001). Food and individual nutrient intakes were calculated using the national food composition database, Fineli (https://fineli.fi/fineli/en/index). We had access to 329 maternal serum samples collected at the beginning of the third trimester and 274 samples taken at delivery. We had, altogether, 300 cord blood samples. By pairing maternal and cord blood samples we obtained 264 paired mother-infant samples.

### DIABIMMUNE study

The DIABIMMUNE study recruited 832 families in Finland (Espoo), Estonia (Tartu), and Russia (Petrozavodsk) with infants carrying HLA alleles that conferred risk for autoimmunity. The subjects involved in the current study were chosen from the subset (n = 52) of international DIABIMMUNE study children who progressed to multiple islet autoantibodies (n = 14) and controls (CTR, n = 38) who remained AAb negative in a longitudinal series of samples collected at 3, 6, 12, 18, 24 and 36 months from each child (Kostic et al., 2015). The study groups were matched by HLA-associated diabetes risk, sex, country and period of birth. This study was conducted according to the guidelines in the Declaration of Helsinki. The Ethics and Research Committee of the participating Universities and Hospitals approved the study protocol. All families provided written informed consent prior to sample collection.

### NOD mouse study – summary

The study setting of the two NOD mouse studies mice was reported previously (Berntsen et al., 2018; Bodin et al., 2016). In short, NOD/ShiLtJ mice from the Jackson Laboratory (Maine, USA) were used for breeding at 8 and 10 weeks of age and randomly allocated to the exposure groups. Female offspring were, in both studies, exposed at mating, through gestation and early life until 11-12 weeks of age when the serum samples were collected, with 4-5 mice kept per cage and 5-8 mice per exposure group.

The exposure in the first study (n=5 per exposure group) was to a mixture of persistent organic pollutants in feed, with a high and a low dose mixture (chemical composition based on human intake (Berntsen et al., 2017; Berntsen et al., 2018)). The total intake of PFAS was 0.14 μg/day and 2.8 μg/day in the (1) low and (2) high dose groups, respectively, corresponding to (1) 1-50 times human serum levels of PFAS and (2) 20-1000 times the human serum levels (Berntsen et al., 2017). In both studies, the mice had *ad libitum* access to food and water (Harlan Teklad 2919 irradiated, Madison, WI) and had a 12 h light/12 h dark cycle with 35–75% humidity. The exposure in the second study was to PFUnDA in the drinking water (n=8 per group) (0, 3, 30 and 300 ▯g/L, corresponding to 0.417, 4.17 and 41.7 μg/kg bw/day). The lowest exposure level of PFUnDA is about five times higher than the maximal calculated intake of PFOA in human infants.

All experiments were performed in conformity with the laws and regulations for experiments with live animals in Norway and were approved by the local representative of the Norwegian Animal Research Authority. In the NOD mouse model, insulitis is the most prominent feature preceding diabetes onset, with impaired macrophage phagocytosis being associated with seroconversion. Insulitis was assessed by grading of hematoxylin and eosin-stained pancreatic tissue sections. Early signs of insulitis included an increased number of apoptotic cells, a decreased number of tissue-resident macrophages in pancreatic islets and reduced phagocytic function of macrophages isolated from the peritoneum.

### Analysis of PFAS

Sample preparation and analysis for PFAS was carried out as described previously (Salihovic et al., 2013). In short, 450 μL acetonitrile with 1 % formic acid, and internal standards were added to 150 μL serum and samples subsequently treated with Ostro sample preparation in a 96-well plate for protein precipitation and phospholipid removal. The analysis of PFAS was performed using automated column-switching ultra-performance liquid chromatography-tandem mass spectrometry (UPLC-MS/MS) (Waters, Milford, USA) using an ACQUITY C18 BEH 2.1×100mm×1.7μm column and a gradient with 30% methanol in 2mM NH4Ac water and 2mM NH4Ac in methanol with a flow rate of 0.3 mL/min. Quantitative analysis of the selected analytes was performed using the isotope dilution method; all standards (*i.e.*, internal standards, recovery standards, and native calibration standards) were purchased from Wellington Laboratories (Guelph, Ontario, Canada). The method’s detection limits ranged between 0.02-0.19 ng/mL, depending on the analyte.

### Analysis of bile acids

The bile acids were measured in NOD mice and in human serum as described recently (Salihovic et al., 2019), with some modifications in the sample preparation. 100 μL of acetonitrile and 10 μL of PFAS internal standard mixture (c = 200 ▯g/mL in methanol) and 20 ▯L of BA internal standard mixture (c = 440-670 ▯g/mL in methanol) and 50 μL NOD serum respectively were mixed, the samples were centrifuged and the organic phase was collected, evaporated to dryness after which ^13^C injection standards were added (10 ▯L of 200 ▯g/mL PFAS in methanol) as was 300 ▯L of 2 mM NH4AC in water. For human samples, 20 μL of serum, using the same internal standard mixtures, was filtered through a frit filter plate (96-Well Protein Precipitation Filter Plate, Sigma Aldrich), and the effluent was collected and evaporated to dryness and the residue was dissolved in 20 ▯L of a 40:60 MeOH:H2O v/v mixture containing the same ^13^C-PFAS injection standards. Analyses were performed on an ACQUITY UPLC system coupled to a triple quadrupole mass spectrometer (Waters Corporation, Milford, USA) with an atmospheric electrospray interface operating in negative ion mode. An external calibration with six calibration points (0.5-160 ng/mL), including a solvent blank, was carried out for use in quantitation.

### Analysis of molecular lipids by UHPLC-QTOFMS

Serum samples were randomized and extracted using a modified version of the previously-published Folch procedure (Nygren et al., 2011). In short, 10 μL of 0.9% NaCl and, 120 μL of CHCl3: MeOH (2:1, v/v) containing the internal standards (c = 2.5 μg/mL) was added to 10 μL of each serum sample. The standard solution contained the following compounds: 1,2-diheptadecanoyl-sn-glycero-3-phosphoethanolamine (PE(17:0/17:0)), N-heptadecanoyl-D-erythro-sphingosylphosphorylcholine (SM(d18:1/17:0)), N-heptadecanoyl-D-erythro-sphingosine (Cer(d18:1/17:0)), 1,2-diheptadecanoyl-sn-glycero-3-phosphocholine (PC(17:0/17:0)), 1-heptadecanoyl-2-hydroxy-sn-glycero-3-phosphocholine (LPC(17:0)) and 1-palmitoyl-d31-2-oleoyl-sn-glycero-3-phosphocholine (PC(16:0/d31/18:1)), were purchased from Avanti Polar Lipids, Inc. (Alabaster, AL, USA), and, triheptadecanoylglycerol (TG(17:0/17:0/17:0)) was purchased from Larodan AB (Solna, Sweden). The samples were vortex mixed and incubated on ice for 30 min after which they were centrifuged (9400 × g, 3 min). 60 μL from the lower layer of each sample was then transferred to a glass vial with an insert and 60 μL of CHCl3: MeOH (2:1, v/v) was added to each sample. The samples were stored at −80 °C until analysis.

Calibration curves using 1-hexadecyl-2-(9Z-octadecenoyl)-sn-glycero-3-phosphocholine (PC(16:0e/18:1(9Z))), 1-(1Z-octadecenyl)-2-(9Z-octadecenoyl)-sn-glycero-3-phosphocholine (PC(18:0p/18:1(9Z))), 1-stearoyl-2-hydroxy-sn-glycero-3-phosphocholine (LPC(18:0)), 1-oleoyl-2-hydroxy-sn-glycero-3-phosphocholine (LPC(18:1)), 1-palmitoyl-2-oleoyl-sn-glycero-3-phosphoethanolamine (PE(16:0/18:1)), 1-(1Z-octadecenyl)-2-docosahexaenoyl-sn-glycero-3-phosphocholine (PC(18:0p/22:6)) and 1-stearoyl-2-linoleoyl-sn-glycerol (DG(18:0/18:2)), 1-(9Z-octadecenoyl)-sn-glycero-3-phosphoethanolamine (LPE(18:1)), N-(9Z-octadecenoyl)-sphinganine (Cer(d18:0/18:1(9Z))), 1-hexadecyl-2-(9Z-octadecenoyl)-sn-glycero-3-phosphoethanolamine (PE(16:0/18:1)) from Avanti Polar Lipids, 1-Palmitoyl-2-Hydroxy-sn-Glycero-3-Phosphatidylcholine (LPC(16:0)), 1,2,3 trihexadecanoalglycerol (TG(16:0/16:0/16:0)), 1,2,3-trioctadecanoylglycerol (TG(18:0/18:0/18:)) and 3β-hydroxy-5-cholestene-3-stearate (ChoE(18:0)), 3β-Hydroxy-5-cholestene-3-linoleate (ChoE(18:2)) from Larodan, were prepared to the following concentration levels: 100, 500, 1000, 1500, 2000 and 2500 ng/mL (in CHCl3:MeOH, 2:1, v/v) including 1250 ng/mL of each internal standard.

The samples were analyzed by ultra-high-performance liquid chromatography quadrupole time-of-flight mass spectrometry (UHPLC-QTOFMS). Briefly, the UHPLC system used in this work was a 1290 Infinity II system from Agilent Technologies (Santa Clara, CA, USA). The system was equipped with a multi sampler (maintained at 10 °C), a quaternary solvent manager and a column thermostat (maintained at 50 °C). Injection volume was 1 μL and the separations were performed on an ACQUITY UPLC^®^ BEH C18 column (2.1 mm × 100 mm, particle size 1.7 μm) by Waters (Milford, MA, USA). The mass spectrometer coupled to the UHPLC was a 6545 QTOF from Agilent Technologies interfaced with a dual jet stream electrospray (Ddual ESI) ion source. All analyses were performed in positive ion mode and MassHunter B.06.01 (Agilent Technologies) was used for all data acquisition. Quality control was performed throughout the dataset by including blanks, pure standard samples, extracted standard samples and control serum samples. Relative standard deviations (% RSDs) for peak areas for lipid standards representing each lipid class in the control serum samples (n = 12) and in the pooled serum samples (n = 77) were calculated on average 15.9% and 13.6% (raw variation) in maternal samples and in cord blood samples, respectively. For serum samples from NOD mice, RSD was on average 11.9%. The lipid concentrations in pooled control samples showed % RSDs within accepted analytical limits at averages of 14.7% and 20.4% for the maternal and cord blood serum samples, respectively, and 7.3% for serum samples from NOD mice.

Mass spectrometry data processing was performed using the open source software package MZmine 2.18 (Pluskal et al., 2010). The following steps were applied in this processing: (i) Crop filtering with a m/z range of 350 – 1200 m/z and an RT range of 2.0 to 12 minutes, (ii) Mass detection with a noise level of 750, (iii) Chromatogram builder with a minimum time span of 0.08 min, minimum height of 1000 and a m/z tolerance of 0.006 m/z or 10.0 ppm, (iv) Chromatogram deconvolution using the local minimum search algorithm with a 70% chromatographic threshold, 0.05 min minimum RT range, 5% minimum relative height, 1200 minimum absolute height, a minimum ration of peak top/edge of 1.2 and a peak duration range of 0.08 – 5.0, (v), Isotopic peak grouper with a m/z tolerance of 5.0 ppm, RT tolerance of 0.05 min, maximum charge of 2 and with the most intense isotope set as the representative isotope, (vi) Peak filter with minimum 12 data points, a FWHM between 0.0 and 0.2, tailing factor between 0.45 and 2.22 and asymmetry factor between 0.40 and 2.50, (vii) Join aligner with a m/z tolerance of 0.009 or 10.0 ppm and a weight for of 2, a RT tolerance of 0.1 min and a weight of 1 and with no requirement of charge state or ID and no comparison of isotope pattern, (viii) Peak list row filter with a minimum of 10% of the samples (ix) Gap filling using the same RT and m/z range gap filler algorithm with an m/z tolerance of 0.009 m/z or 11.0 ppm, (x) Identification of lipids using a custom database search with an m/z tolerance of 0.009 m/z or 10.0 ppm and a RT tolerance of 0.1 min, and (xi) Normalization using internal standards PE(17:0/17:0), SM(d18:1/17:0), Cer(d18:1/17:0), LPC(17:0), TG(17:0/17:0/17:0) and PC(16:0/d30/18:1)) for identified lipids and closest ISTD for the unknown lipids followed by calculation of the concentrations based on lipid-class concentration curves.

An aliquot of each sample was collected and pooled and used as quality control sample, together with NIST SRM1950 reference plasma sample, an in-house pooled serum sample.

### Analysis of polar metabolites by GC-QTOFMS

Serum samples were randomized and sample preparation was carried out as described previously (Castillo et al., 2011). In summary, 400 μL of MeOH containing ISTDs (heptadecanoic acid, deuterium-labeled DL-valine, deuterium-labeled succinic acid, and deuterium-labeled glutamic acid, c = 1 μg/mL) was added to 30 μl of the serum samples which were vortex mixed and incubated on ice for 30 min after which they were centrifuged (9400 × g, 3 min) and 350 μL of the supernatant was collected after centrifugation. The solvent was evaporated to dryness and 25 μL of MOX reagent was added and the sample was incubated for 60 min at 45 °C. 25 μL of MSTFA was added and after 60 min incubation at 45 °C 25 μL of the retention index standard mixture (n-alkanes, c=10 μg/mL) was added.

The analyses were carried out on an Agilent 7890B GC coupled to 7200 QTOF MS. Injection volume was 1 μL with 100:1 cold solvent split on PTV at 70 °C, heating to 300 °C at 120 °C/minute. Column: Zebron ZB-SemiVolatiles. Length: 20m, I.D. 0.18mm, film thickness: 0.18 μm. With initial Helium flow 1.2 mL/min, increasing to 2.4 mL/min after 16 mins. Oven temperature program: 50 °C (5 min), then to 270°C at 20 °C/min and then to 300 °C at 40 °C/min (5 min). EI source: 250 °C, 70 eV electron energy, 35μA emission, solvent delay 3 min. Mass range 55 to 650 amu, acquisition rate 5 spectra/s, acquisition time 200 ms/spectrum. Quad at 150 °C, 1.5 mL/min N2 collision flow, aux-2 temperature: 280 °C.

Calibration curves were constructed using alanine, citric acid, fumaric acid, glutamic acid, glycine, lactic acid, malic acid, 2-hydroxybutyric acid, 3-hydroxybutyric acid, linoleic acid, oleic acid, palmitic acid, stearic acid, cholesterol, fructose, glutamine, indole-3-propionic acid, isoleucine, leucine, proline, succinic acid, valine, asparagine, aspartic acid, arachidonic acid, glycerol-3-phosphate, lysine, methionine, ornithine, phenylalanine, serine and threonine purchased from Sigma-Aldrich (St. Louis, MO, USA) at concentration range of 0.1 to 80 μg/mL. An aliquot of each sample was collected and pooled and used as quality control samples, together with a NIST SRM 1950 serum sample and an in-house pooled serum sample. Relative standard deviations (% RSDs) of the metabolite concentrations in control serum samples showed % RSDs within accepted analytical limits at averages of 12.3% and 19.6% for the maternal and cord blood serum samples, respectively, and 7.2% for serum samples from NOD mice.

### Analysis of islet autoantibodies (EDIA and DIABIMMUNE studies)

Four diabetes-associated autoantibodies were analyzed from each serum sample with specific radiobinding assays: insulin autoantibodies (IAA), glutamic acid decarboxylase antibodies (GADA), islet antigen-2 antibodies (IA-2A), and zinc transporter 8 antibodies (ZnT8A) as described previously (Kostic et al., 2015). Islet cell antibodies (ICA) were analyzed with immunofluoresence in those subjects who tested positive for at least one of the biochemical autoantibodies. The cut-off values were based on the 99th percentile in non-diabetic children and were 2.80 relative units (RU) for IAA, 5.36 RU for GADA, 0.78 RU for IA-2A and 0.61 RU for ZnT8A. The detection limit in the ICA assay was 2.5 Juvenile Diabetes Foundation units (JDFU).

### Statistical analysis

All analyses here were performed using the R statistical programming language (https://www.r-project.org/).

For all datasets, the following preprocessing steps were carried out:

1. NA or negative values in the data were replaced with zeroes.
2. All zeroes were then replaced with imputed half-minimums (for each variable, the minimum value was found, and half of this value was used).
3. All values were log2 transformed.
4. Each variable was scaled to zero mean and unit variance.

#### Clustering of metabolites

All metabolomics datasets were then analyzed using the *mclust* R package (version 5.4.1) to assign variables (lipids / metabolites) from each dataset to separate clusters. Here, *mclust* attempts to fit various model types and assesses their performance using the Bayesian Information Criterion (BIC). Maximization of the BIC is a well-established method for model selection, particularly useful in the case of clustering of data, to choose the optimum number of clusters by way of maximizing the likelihood of fit of the clustering model, whilst this metric penalizes (and thereby avoids) unnecessary complexity. The BIC is calculated at each iteration, and the optimal (maximal) BIC will occur when the lowest number of clusters are used before the point at which increasing the number of clusters gives a lesser return in the fitness of the model. The highest BIC achieved by *mclust* for each dataset was therefore used to determine both the model type and the number of clusters into which the variables should be divided. The variables in each dataset were accordingly given numbers to denote their cluster membership.

For each sample in each dataset, cluster variables were then generated. For each dataset, the number of clusters *k* found by *mclust* equals the number of cluster variables generated. Each cluster variable is the mean value of the lipids / metabolites that make up that cluster, meaning that samples in that dataset become represented only by (n = *k)* values. These cluster variables were given acronyms indicating the dataset from which they were generated (Maternal blood Lipids Cluster – MLC, Maternal polar Metabolites Cluster - MMC, Child Lipids Cluster - CLC, Child polar Metabolites Cluster – CMC).

These cluster variable acronyms had their cluster numbers (1-*k* for each dataset) appended to them to form their final labels. Assignment of individual lipids / metabolites to each cluster variable is given for each dataset in supplementary files.

#### Correlation analysis

With dataset dimensionality reduced to the aforementioned cluster variables, correlation analysis was employed, taking all cluster variables into account, along with clinical variables, simultaneously. Spearman correlations were calculated pairwise between all of the aforementioned variables. To subsequently visualize this, the *corrplot* R package (version 0.84) was used. For legibility of figures, the colors of these plots generated by *corrplot* were limited to either solid orange or blue for positive or inverse correlations respectively, with correlation strength represented purely through the size of the filled circles for each pairwise correlation.

#### Partial correlation network

For network analysis and representation based on the generated partial correlations, the *qpNrr* function from the *qpgraph* R package (version 2.16.0) was run with default parameters to estimate non-rejection rates (NRRs) of the aforementioned correlation matrix of all datasets’ cluster variables and clinical features. This is a means for rejecting spurious correlations. The obtained NRR matrix was then filtered at various thresholds (0.1 to 1, with an increment of 0.1) to select the edges included in network representation. The distribution of NRRs was also visualized as a histogram to assist with choosing an appropriate cut-off threshold for retaining plausible, and rejecting likely spurious, correlations. Based on the distribution of NRRs and observed network topology of the generated networks, a conservative cut-off of 0.4 was deemed appropriate.

The *Rgraphviz* R package (version 2.26.0) was used to generate network topography plots. Node and edge properties for these network graphs were generated in a custom fashion. Edges were colored by the directionality of the relationship between the nodes that they connect. Edge width was plotted as a function of the strength of the Spearman correlation between the two variables that the edge connects. Nodes were colored and shaped purely for clarity and to group like variables (PFAS, clinical variables, cluster variables) and sample sources (mothers, infants) together. Network layout is generated by the *Rgraphviz* package itself, and layout was set to the “neato” parameter to balance clarity and compactness.

For both the final correlation plot and network figure, values were used only from those sample identifiers uniquely represented in all involved datasets (maternal blood lipids, maternal polar metabolites, child blood lipids, child blood polar metabolites, HLA data, demographic data), totaling (n = 226 samples, n = 54 features).

#### Univariate statistical methods

To identify any child lipid / metabolite clusters (CLCs and CMCs) changing in value across various levels of maternal PFAS exposure, the mothers were first assigned to four quartiles of total PFAS exposure, defined as the sum of their raw PFAS exposures. Subsequently, an analysis of variance (ANOVA) test was performed to assess any significant change in the means of the CLCs / CMCs across maternal PFAS exposure quartiles. To further identify the specific quartiles between which any such significant changes in the CLCs / CMCs took place, Tukey’s honest significant differences (Tukey HSDs) were calculated with the same design, enabling intra-PFAS exposure quartile comparisons of CLCs / CMCs.

Multi-way analysis of variance was performed with factors HLA risk and PFAS exposure) and their interactions in MATLAB R2017b (Mathworks, Inc., Natick, MA, USA) using the Statistical Toolbox.

The Wilcoxon rank-sum test was used in comparing the two study groups of samples (*e.g.* CTR *vs.* mAAb+ group) in a specific age cohort. These statistical analyses were computed in MATLAB 2017b using the statistical toolbox. For statistical comparison, subjects with missing peaks for the given quantified compound and children who were not exclusively breast-fed for 30 days were excluded. The longitudinal profiles of the PFOS in the samples obtained from children who progressed to multiple autoantibodies and autoantibody negative controls were compared using linear mixed-effects model with the fixed effect being case, age, sex, and the random effect being subject-wise variation using lme4 package in R. The fully-parametrized model was compared with a null model using analysis of variance (ANOVA) as deployed in the *lme4* R package. The locally-weighted regression plot was made using smoothing interpolation function loess (with span = 0.85) available from *ggplot2* package in R. The individual metabolite levels were visualized as scatter plot as well as box plot using GraphPad Prism 7 (GraphPad Software, La Jolla, CA, USA).

#### Regression analysis

Linear regression (LR) with L2 regularization was performed to assess the effect of individual maternal PFAS on the significantly altered (ANOVA, Tukey’s HSD test, p < 0.05) cord blood lipids, polar metabolites and their cluster, between the higher (Q4) and the lower (Q1) exposure level. Concentrations of 20 measured PFAS were regressed against the metabolite intensities or their mean cluster profiles. Regularized LR modelling was performed using *‘glmnet’* function deployed in the R-package *‘glmnet v2.0-18’.* The hyper-parameter ‘λ_minimum_’ was selected based on the minimum cross-validation (CV) error as determined by 10-fold CV using *‘cv.glmnet’*. The predictors were standardized. The LR models were adjusted for covariates such as mothers’ age and BMI. In order to get the ranks of the individual predictor, ridge-coefficients were estimated and normalized with the maximum value. In addition, iterative step-wise LR models were developed for the selection of minimum number of model variables that are required to maximize the outcome. The step-wise LR models with the minimum Akaike information criterion (AIC) was considered. The top predictors/variables of the ridge models were also selected by the step-wise LR models.

Further, to quantify the effect of individual or combination of PFAS as regard to the stratification of AAb+ *vs.* AAb-groups in a follow-up EDIA study, we performed an iterative step-wise approach to develop predictive logistic ridge regression (LRR) models. The regularization strategy was adapted to avoid any issues concerning multicollinearity among the highly-correlated predictors. The LRR models were bootstrapped, where 80% of the mothers’ PFAS exposure samples were used to train the model whilst 20% was used as a test data. In order to make AAb-group (n>>10) comparable with AAb+ group (n=10), downsampling was performed. This was repeated 10,000 times. The partitioning of the data was achieved by the *‘caret 6.0.84’* R package. The model with the highest mean AUC was considered to be the best model as assessed by their receiver operating characteristic (ROC) curve and 95% CI *via* the *‘pROC 1.15.3’ R* package. All the LRR models with AUC > 0.60 (with 10-fold cross-validation) were considered. The odds ratio and CI for each predictor was estimated. LRR modelling was performed using R-package *‘glmnet v2.0-18’.* The hyper-parameter ‘λ_minimum_’ was selected based on 10-fold CV. The recursive step-wise feature elimination scheme guided in the optimal selection of the PFAS that aids in the separation of AAb+ *vs.* AAb-groups. The PFAS in LRR models were either incorporated or removed in an iterative manner, starting with all 20 PFAS. LRR models were adjusted for mothers’ age and BMI. Accuracy of prediction was determined by AUCs.

Further, we performed LRR modelling to determine the ranks of 20 PFAS in separation of AAb+ vs. AAb-. The ranks of the predictors (PFAS) were estimated as the unit absolute differences in the odd ratios, *i.e.* predictors with maximum absolute differences were ranked as the top contributors. The hyper-parameter ‘λ_minimum_’ for LRR was selected based on 10-fold CV.

## Acknowledgments

The authors thank Samira Salihović and Tuomas Lindeman for technical expertise in the development of bile acid analysis method. Bile acid analyses were partly performed at the Turku Metabolomics Centre, core facility that is part of the Biocenter Finland national research infrastructure.

The EDIA study was supported by the National Institute of Diabetes and Digestive and Kidney Diseases (NIDDK), National Institutes of Health (No. 1DP3DK094338-01 to M.K.), the Academy of Finland Centre of Excellence in Molecular Systems Immunology and Physiology Research 2012-17, No. 250114 to M.K. and M.O.), Academy of Finland postdoctoral grant (No. 323171 to S.L.) and the Medical Research Funds, Tampere and Helsinki University Hospitals (to M.K.). The current work was also supported by the Swedish Research Council (No. 2016-05176 to T.H.), Formas (No. 2019-00869 to T.H. and M.O.), Juvenile Diabetes Research Foundation (2-SRA-2016-341-S-B to M.K., T.H. and M.O., 2-SRA-2016-289-S-B to M.O.), Academy of Finland (No. 292568 to M.O.), and the Knowledge Foundation (to T.H.).

## Author contributions

T.H. initially proposed the study of PFAS exposure in the mother-infant study (EDIA), while M.O. proposed the follow-up metabolomics studies in NOD mice. T.H., M.O. and M.K. initiated and designed the study. M.K. is the principal investigator of the mother-infant study. M.O. supervised data analysis and integration, and together with T.H. did the primary interpretation of analytical outcomes. A.M., M.O., S.L. P.S. and T.H. analyzed the data. T.H. supervised metabolomic and PFAS analyses. T.S., D.G., C.C., D.D. and A.D. performed sample analyses. J.B., H.D. and U.C.N. conducted the studies in NOD mice, while H.F.B. and K.Z. assembled the internal relationship of chemicals in the feed for POP-exposure. H.S. contributed clinical research in the mother-infant cohort. J.I. performed genetic analyses in the mother-infant study. S.M.V. performed the study of dietary intake in the mother-infant study. A.M., M.O. and T.H. wrote the first version of the paper. All authors approved the final version.

## Supplementary Materials

**Fig. S1.**
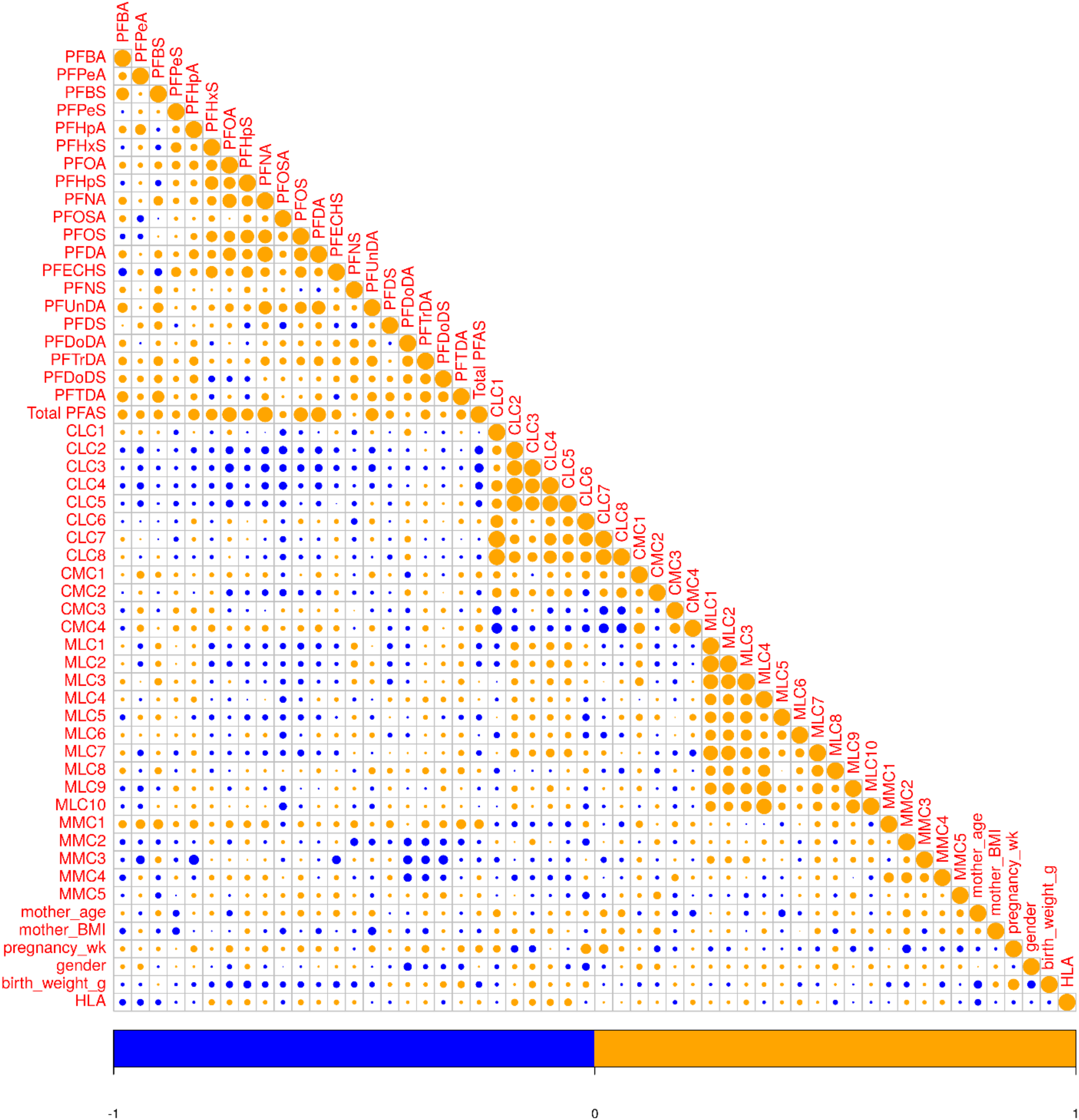
Correlation plot of PFAS exposure, metabolite clusters and demographic data in the mother-infant cohort. Pairwise Spearman correlation as calculated between all cluster variables, PFAS, demographic variables and HLA status in the mother-child cohort. Positive correlations in orange, negative correlations in blue. Dot size for each pairwise correlation corresponds to the strength of the calculated correlation.

**Fig. S2.**
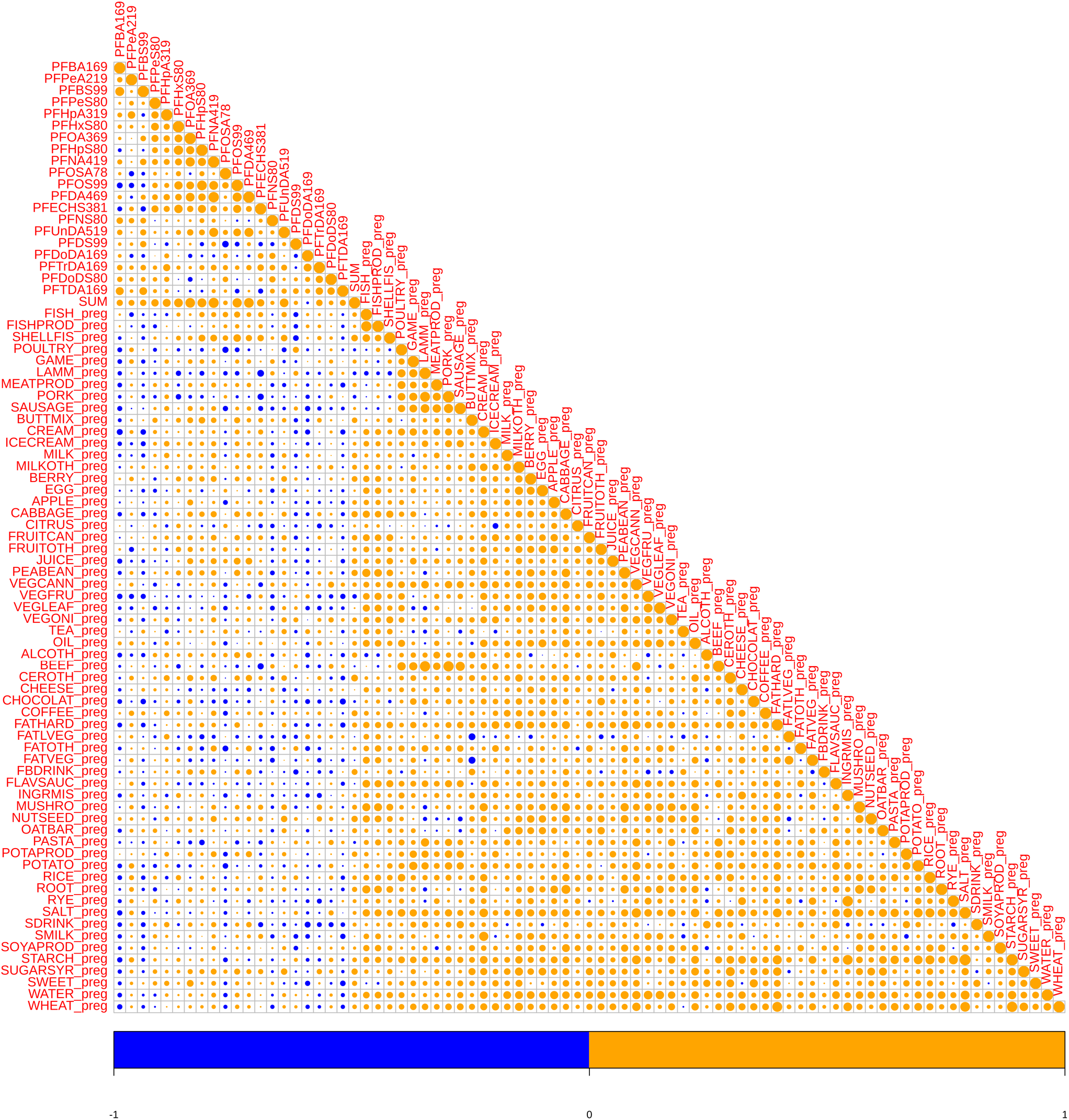
Correlation of PFAS exposure and dietary factors during pregnancy. Pairwise Spearman correlation as calculated between all mothers’ measured serum PFAS concentrations and dietary variables. Positive correlations in orange, negative correlations in blue. Dot size for each pairwise correlation corresponds to the strength of the calculater correlation.

**Fig. S3.**
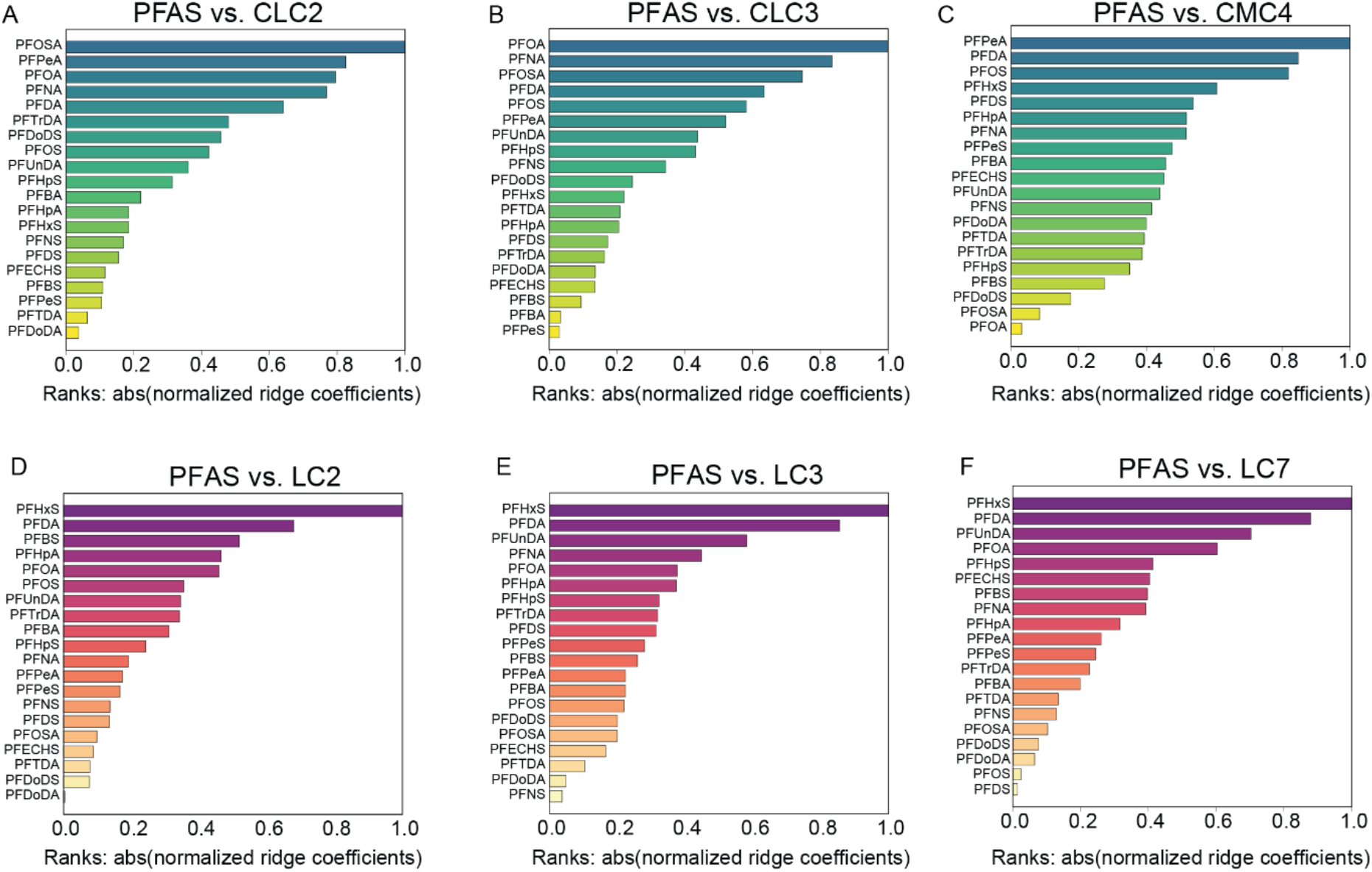
Identification of the most contributing PFAS to selected metabolite/lipid clusters. (A-C) Bar plots showing mothers’ PFAS exposures as linear predictors of metabolic change in selected cord serum lipid/metabolite clusters (CLC, CMC). The PFAS are ranked and sorted by their regression (ridge) coefficients, *i.e.*, the most contributing PFAS (predictor) associated with the clusters are shown at the top of the chart. These selected lipid/metabolite clusters were significantly (adjusted p < 0.05) altered between highest (Q4) and the lowest (Q1) quartiles of the total maternal PFAS exposure. (D-F) Showing mothers’ PFAS exposures as predictors of T1D lipid clusters (LCs) of the newborn infants, based on LCs derived in an earlier study (Oresic et al., 2013).

**Fig. S4.**
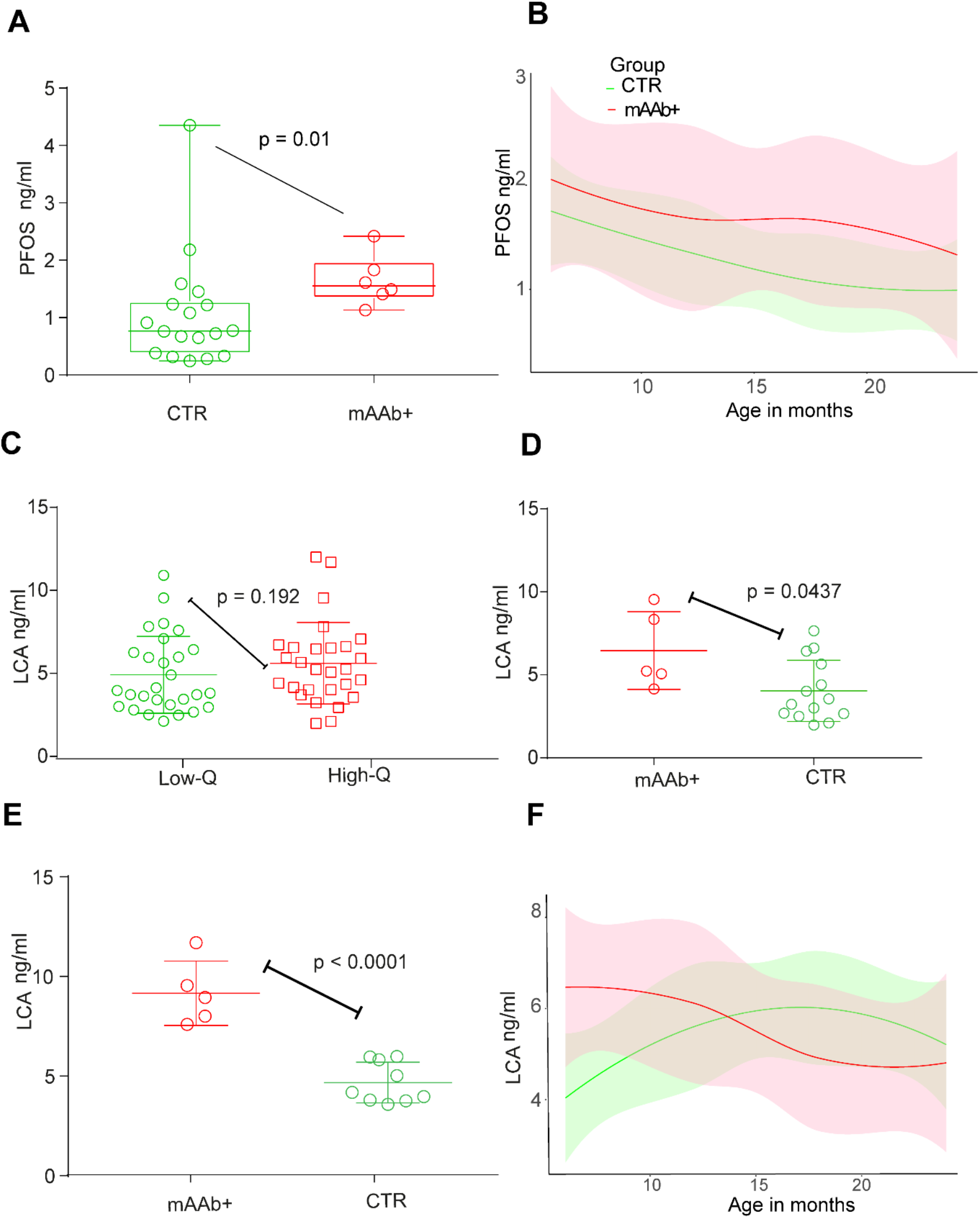
Comparison of PFOS and LCA concentrations between control children and children who progressed to multiple islet autoantibodies during follow-up in the DIABIMMUNE study. (A) Concentration of PFOS at 18 months of age. Here, CTR; n = 18, mAAb+; n = 6. (B) The loess curve plot of PFOS over time (6, 12, 18, and 24) between the CTR and mAAb+ groups. (C) Concentrations of LCA between children with the lowest level (Q1) of PFOS exposure (Low Q) and those with highest level (Q4) of PFOS exposure (High Q). (D) Scatter plot comparing levels of LCA between control children (CTR) and those who progressed to multiple islet autoantibodies during follow-up (mAAb+) at the age of 6 months (CTR; n = 14, mAAb+; n = 5) (E) and 36 months (CTR; n =9, mAAb+; n = 5) (F) The loess curve plot of LCA over time (6, 12, 18, and 24) between CTR and two Ab group. Only children who were breastfed (≥ 30 days) were included in the comparison.

**Table S1.**
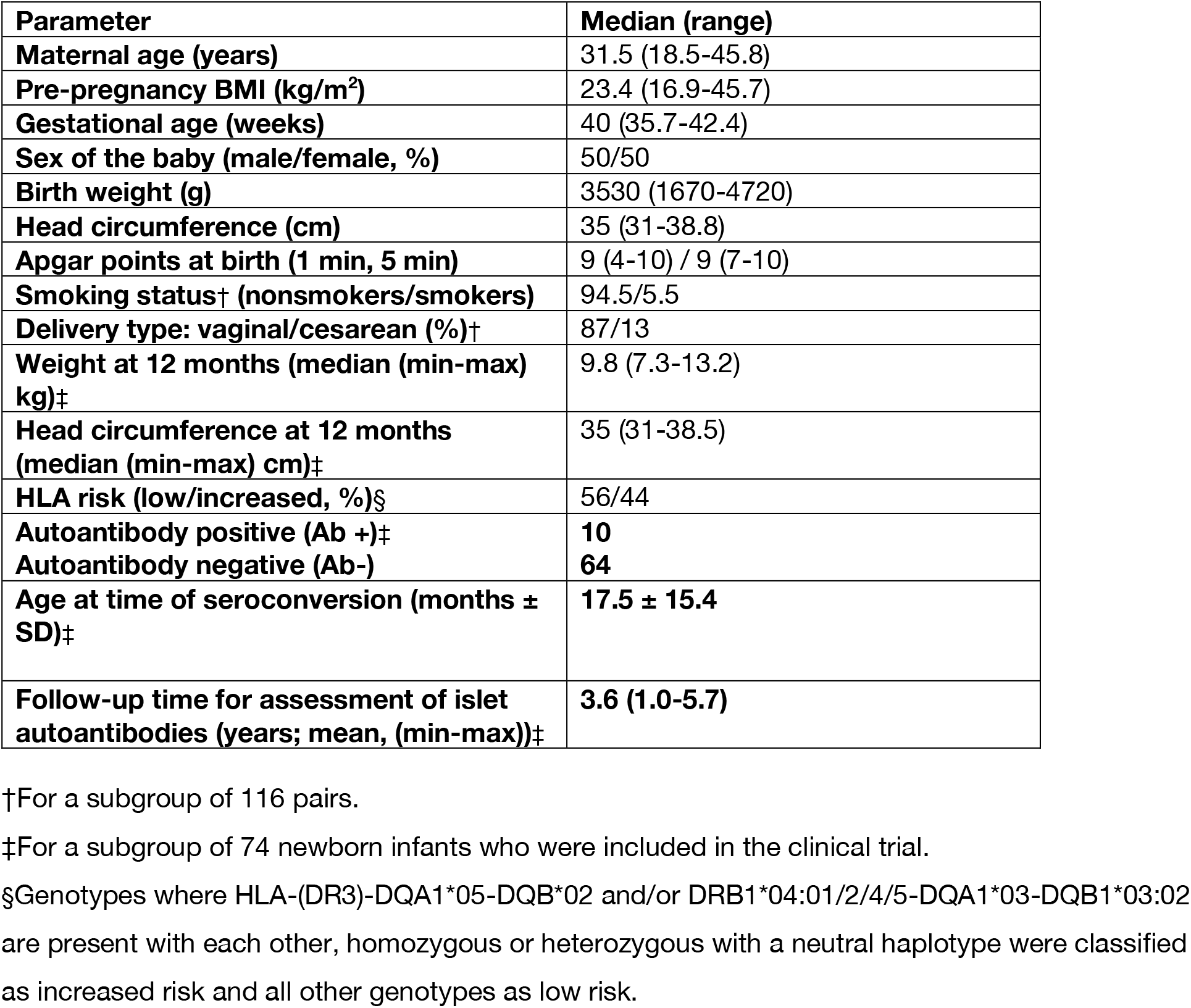
Demographic characteristics of the study population (mother-child cohort).

**Table S2.**
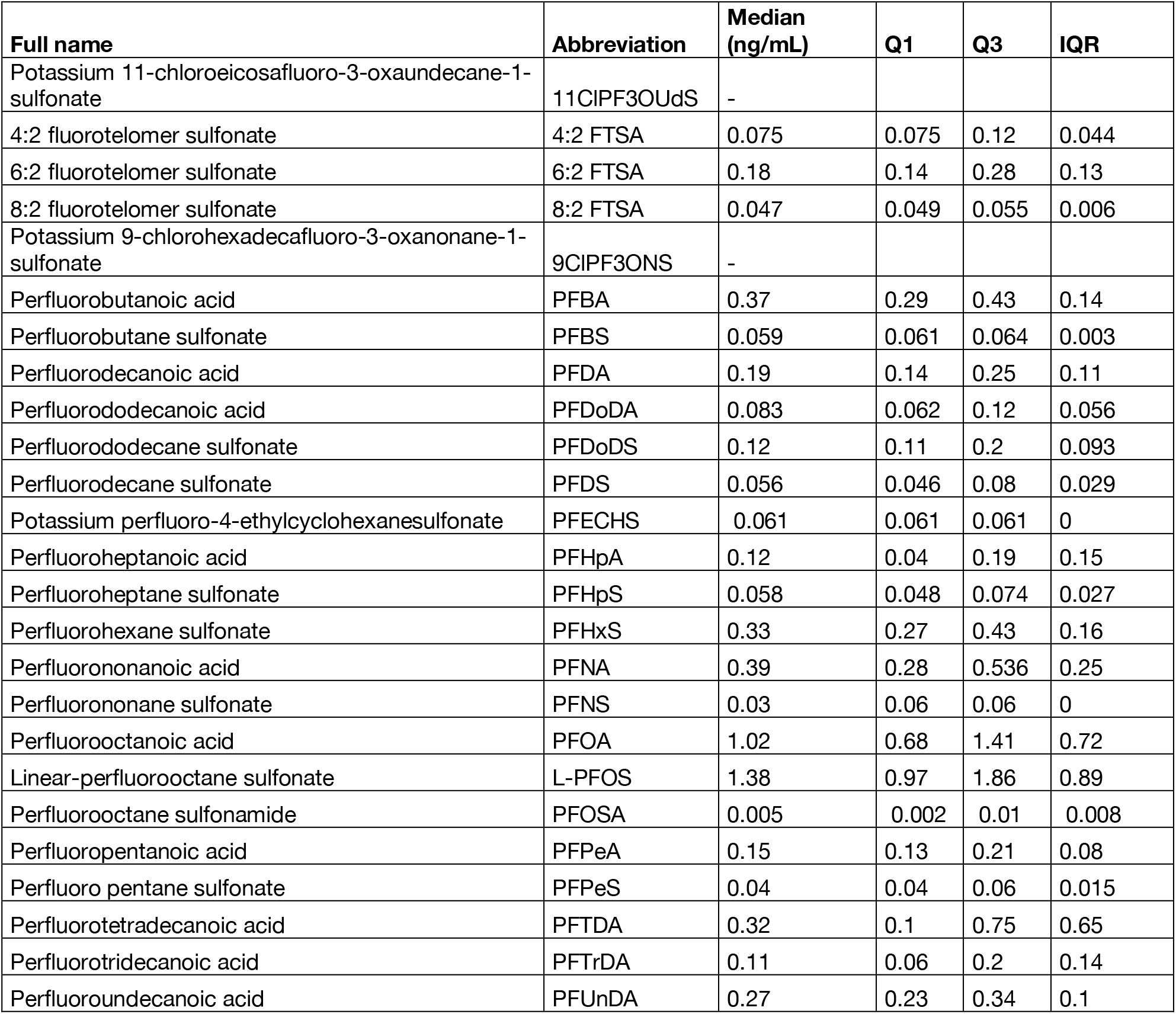
List of PFAS measured in the pregnant mothers. Abbreviations: Q, quartile; IQR, inter-quartile range.

**Table S3.** Description of the lipid and polar metabolite clusters from mothers and offspring. Abbreviations: BCAA, branched chain amino acid; CE, cholesterol ester; LPC, lysophosphatidylcholine, PC, phosphatidylcholine; PE, phosphatidylethanolamine; PUFA, polyunsaturated fatty acids; SFA, saturated fatty acics; TG, triacylglycerol.

| Cluster | Description | Most abundant lipids |
| --- | --- | --- |
| MLC1 | Major phospholipids, one CE | CE(16:0), PC(34:2), PC(16:0/18:1), PC(36:2) |
| MLC2 | Phospholipids, cholesterol esters | CE(18:1), PC(32:1), PC(36:3), PC(36:5) |
| MLC3 | Ceramides | Cer(d18:1/24:1), Cer(d18:1/24:0) |
| MLC4 | TGs with C50-52, mainly unsaturated, mono-unsaturated | TG(18:1/18:1/16:0), TG(18:2/18:1/16:0)<br>TG(14:0/18:1/18:1) |
| MLC5 | LPCs | LPC(18:0), LPC(18:1), LPC(18:2) |
| MLC6 | Ether phospholipids | PE(O-16:0/22:6), PC(O-38:5) |
| MLC7 | PEs | PE(38:6), PE(38:4), PE(16:0/18:1) |
| MLC8 | Small TGs (C<50) with SFA. | TG(14:0/16:0/18:1), TG(18:1/12:0/18:1), TG(16:0/16:0/16:0) |
| MLC9 | Large TGs (>54) with PUFAs | TG(18:2/22:5/16:0), TG(16:0/22:5/18:1), TG(54:6) |
| MLC10 | Medium TGs with PUFA | TG(18:1/18:1/18:1), TG(18:2/18:1/18:1)<br>TG(18:1/18:2/18:2) |
| MMC1 | Amino acids, free cholesterol, saturated free fatty acids | Alanine, decanoic acid, lactic acid, octanoic acid, palmitic acid |
| MMC2 | Amino acids and hydroxy acids | Glutamic acid, glycine, serine |
| MMC3 | Main free fatty acids, amino acids | Glutamine, linoleic acid, oleic acid, stearic acid |
| MMC4 | Branched-chain amino acids | Isoleucine, leucine, valine |
| MMC5 | Various metabolites | 2- and 3-hydroxybutyric acids, indole-3-propionic acid |
| CLC1 | Abundant TGs, mainly containing MUFAs | TG(18:2/18:1/16:0), TG(18:2/18:1/18:1), TG(18:1/18:1/18:1) |
| CLC2 | Abundant PCs and SMs | SM(38:1), SM(42:2), PC(34:2), PC(36:4) |
| CLC3 | LPCs | LPC(18:0), LPC(18:1), LPC(20:4) |
| CLC4 | Phospholipids with PUFAs | PC(36:5), PC(40:7), PE(38:6) |
| CLC5 | Odd-chain PCs | PC(35:1), PC(33:1), PC(35:2) |
| CLC6 | Odd-chain TGs | TG(45:0), TG(47:0), TG(47:1), TG(47:2) |
| CLC7 | Abundant TGs, mainly containing PUFAs | TG(18:0/18:1/20:4), TG(54:6), TG(58:9) |
| CLC8 | Various CE, PC and TG. | CE(20:4), PC(40:7), TG(60:7) |
| CMC1 | Amino acids, dicarboxylic and hydroxyl acids | Alanine, glycine, glutamic acid, fumaric acid, malic acid, citric acid, lactic acid |
| CMC2 | 2- and 3-hydroxybutyric acids, free fatty acids | 2-hydroxybutyric acid, 3-hydroxybutyric acid, palmitic acid, linoleic acid, oleic acid, stearic acid |
| CMC3 | BCAAs, sugar derivatives | Valine, leucine, isoleucine, indole-3-propionic acid |
| CMC4 | Amino acids | Serine, methionine, aspartic acid, phenylalanine |

**Table S4.**
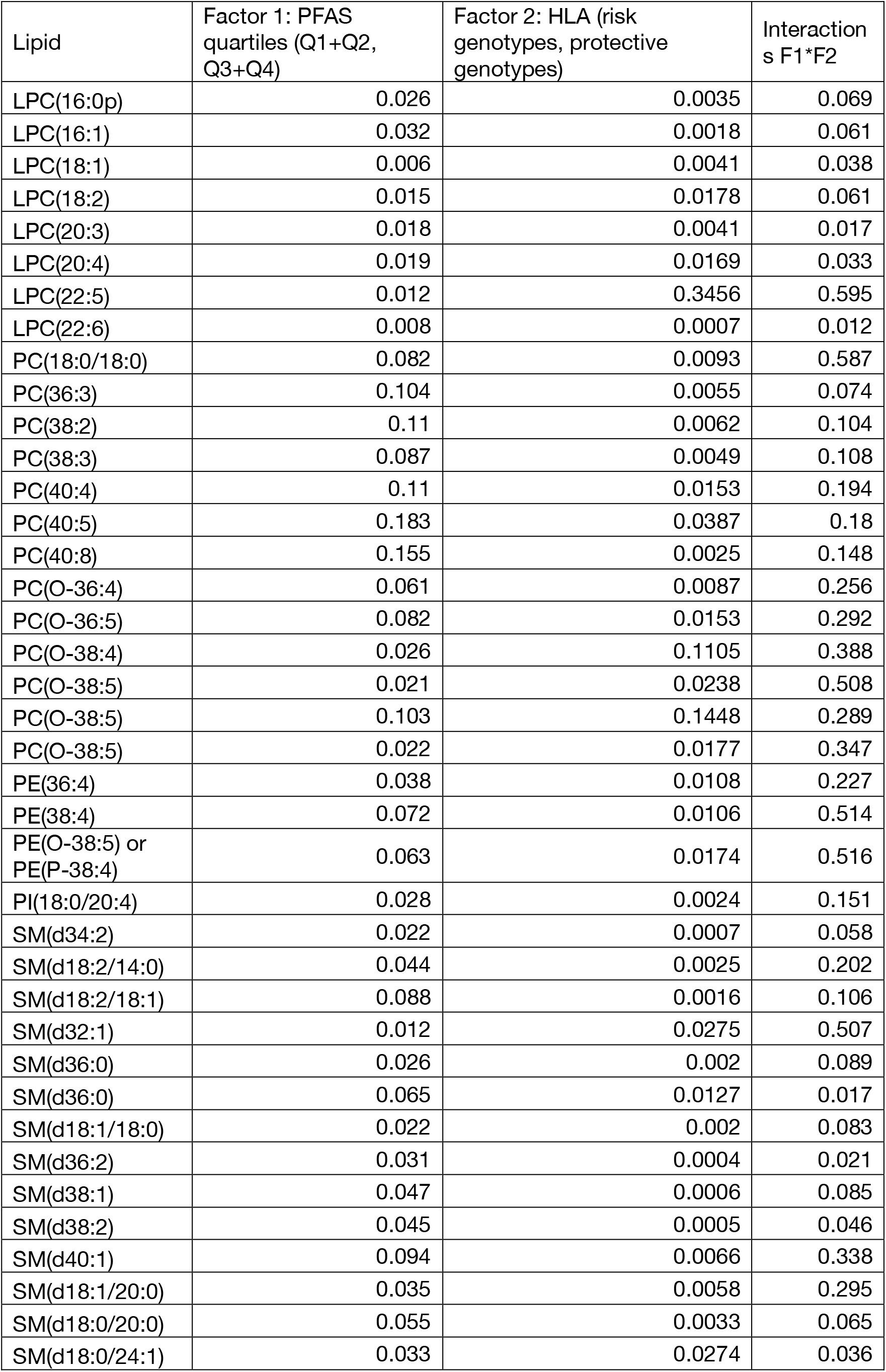
Two-way analysis of variance for cord serum lipids, affected by PFAS exposure and HLA-associated T1D risk. Samples are assigned according to HLA-associated T1D risk (strongly protective / neutral *vs*. mild / high risk) and prenatal total PFAS exposure (quartiles 1 & 2 *vs*. 3 & 4). p-values are shown.

**Table S5.** Bile acids in NOD mice exposed to POP mixture at two different exposure levels. p-values shown (q < 0.05 marked bold, q-values < 0.1 in italics).

|  | Fold<br>(Low/Ctrl) | p (Low/Ctrl) | Fold<br>(High/Ctrl) | p<br>(High/Ctrl) |
| --- | --- | --- | --- | --- |
| TabMCA | 3.57 | 0.140 | 0.79 | 0.254 |
| 7-oxo-DCA | 2.28 | 0.505 | 0.48 | 0.356 |
| 7-oxo-HCA | 4.02 | 0.538 | 0.20 | **0.012** |
| bMCA | 2.61 | 0.279 | 0.17 | **0.006** |
| CA | 1.05 | 0.973 | 0.18 | 0.117 |
| DCA | 1.97 | 0.309 | 0.28 | **4.34×10<sup>-4</sup>** |
| GCA | 2.77 | 0.149 | 0.74 | 0.186 |
| GCDCA | 0.99 | 0.891 | 0.91 | 0.227 |
| GDCA | 1.16 | 0.862 | **0.09** | **2.46×10<sup>-4</sup>** |
| GLCA | 0.91 | 0.506 | 0.90 | 0.302 |
| GUDCA | 0.64 | 0.809 | 0.59 | 0.548 |
| HDCA | 1.38 | 0.196 | **0.23** | **3.79×10<sup>-3</sup>** |
| LCA | **2.14** | **1.15×10<sup>-4</sup>** | **5.92** | **8.90×10<sup>-10</sup>** |
| TCA | 6.28 | 0.090 | 2.23 | 0.816 |
| TCDCA | 3.17 | 0.109 | 0.70 | 0.267 |
| TDCA | 1.55 | 0.686 | **0.05** | **2.41×10<sup>-6</sup>** |
| THDCA | 3.32 | 0.234 | 0.27 | 0.235 |
| TLCA | 1.41 | 0.228 | 0.66 | 0.873 |
| TUDCA | 3.35 | 0.282 | 0.58 | 0.050 |
| TwMCA | 6.79 | 0.110 | 1.01 | 0.442 |
| waMCA | 4.85 | 0.664 | 0.11 | 0.230 |

